# Inhibitory control of synaptic signals preceding motor action in mouse frontal cortex

**DOI:** 10.1101/2021.07.05.451151

**Authors:** Chun-Lei Zhang, Fani Koukouli, Manuela Allegra, Cantin Ortiz, Hsin-Lun Kao, Uwe Maskos, Jean-Pierre Changeux, Christoph Schmidt-Hieber

## Abstract

Preparatory activity in the frontal cortex preceding movement onset is thought to represent a neuronal signature of motor planning. However, how excitatory and inhibitory synaptic inputs to frontal neurons are integrated during movement preparation remains unclear. Here we address this question by performing *in vivo* whole-cell patch-clamp recordings in the secondary motor cortex (MOs) of head-fixed mice moving on a treadmill. We find that both superficial and deep principal neurons show slowly increasing (~10 s) membrane potential and spike rate ramps preceding the onset of spontaneous, self-paced running periods. By contrast, in animals trained to perform a goal-directed task, both membrane potential and spike ramps are characterized by larger amplitudes and accelerated kinetics during preparation of goal-driven movement. To determine the role of local inhibitory neurons in shaping these task-dependent preparatory signals, we chemogenetically suppressed the activity of specific interneuron subpopulations in untrained animals. Inactivation of parvalbumin-positive (PV+) interneurons leads to depolarized membrane potential ramps with increased amplitudes during preparation of movement, while inactivation of somatostatin-positive (SOM+) interneurons abolishes membrane potential ramps. A computational model of the local MOs circuit shows that SOM+-mediated inhibition of PV+ interneurons in conjunction with recurrent connectivity among the principal neurons can reproduce slow ramping signals, while plasticity of excitatory synapses on SOM+ interneurons can explain the acceleration of these signals in trained animals. Together, our data reveal that local inhibitory neurons play distinct roles in controlling task-dependent preparatory ramping signals when MOs neurons integrate external inputs during motor planning.

**Highlights:**

- Principal neurons in MOs show slow preparatory membrane potential and firing rate ramps preceding the onset of spontaneous, self-paced running periods.
- In animals trained to perform a goal-directed task, both membrane potential and spike ramps are faster and larger in amplitude.
- Inactivation of PV+ interneurons disinhibits MOs principal neurons and increases the amplitude of membrane potential ramps, while inactivation of SOM+ interneurons abolishes membrane potential ramps.
- Our modeling results suggest that the concerted action of external inputs and local inactivation shapes task-dependent preparatory motor signals in MOs neurons.

## INTRODUCTION

Neurons in several fronto-parietal brain regions of rodents and primates show gradually increasing ramps of spiking activity that reach a threshold level just before the onset of movement. This preparatory activity has been interpreted as an early neural correlate of motor planning that instructs future actions (Chen et al., 2017; Hanes and Schall, 1996; Inagaki et al., 2019; Li et al., 2015; Maimon and Assad, 2006; Quintana and Fuster, 1999; Roitman and Shadlen, 2002; Thura and Cisek, 2014). It can occur concurrently in multiple areas within the fronto-parietal cortices (Erlich et al., 2015; Goard et al., 2016); alternatively, it appears in a restricted anterior-lateral motor region of the rodent frontal cortex, from which it then spreads to other areas (Chen et al., 2017; Guo et al., 2014). Interactions between several brain regions, including thalamus and cerebellum, are involved in producing and maintaining this ramping neural activity in frontal cortices (Dacre et al., 2021; Gao et al., 2018; Goldman-Rakic, 1995; Tanaka, 2007). In particular, coordinated neural processing in a cortico-thalamic-cerebellar loop is required to maintain preparatory activity across brain regions and to enable successful motor planning (Chabrol et al., 2019; Gao et al., 2018; Wagner et al., 2019).

What is the synaptic basis of this preparatory activity? While the circuit mechanisms underlying these neuronal dynamics in motor-associated cortices have been extensively studied, it is unclear how excitatory and inhibitory synaptic inputs are integrated by individual neurons during preparation of motor actions (Verduzco-Flores et al., 2009). The frontal cortex is characterized by a layered structure containing numerous types of excitatory and inhibitory neurons (DeFelipe and Fariñas, 1992; Kawaguchi and Kubota, 1997). Parvalbumin-positive (PV+) and somatostatin-positive (SOM+) interneurons are two principal subtypes of cortical GABAergic neurons that differ in morphology, physiological properties and targeting of principal neurons (Hangya et al., 2014; Hu et al., 2014; Rudy et al., 2011). Recent intracellular recordings have provided evidence that the interplay between inhibition from these interneurons, excitatory synaptic inputs, and intrinsic membrane properties governs the subthreshold membrane potential dynamics during different locomotor states in several neocortical regions (Gentet et al., 2010; Polack et al., 2013; Schneider et al., 2014). These findings point towards a critical role for synaptic integration of excitatory and inhibitory inputs in shaping preparatory signals in the frontal cortex.

Among the frontal cortical regions that display pre-movement ramping activity, the secondary motor cortex (MOs, or M2) is thought to be particularly important for motor planning (Barthas and Kwan, 2017; Erlich et al., 2011; Murakami et al., 2014). It receives inputs from numerous sensory cortical and thalamic sources, and projects along the corticospinal tract to the spinal cord and superior colliculus to drive movement output (Donoghue and Wise, 1982; Gabbott et al., 2005). Due to its remarkable connectivity, it has been proposed that MOs integrates multi-sensory inputs and organizes motor output during voluntary motion (Barthas and Kwan, 2017; Coen et al., 2021). Here we sought to identify how synaptic inputs are integrated during movement preparation by performing *in vivo* whole-cell patch-clamp recordings from MOs principal neurons in awake mice during resting and running on a treadmill. We find that MOs neurons exhibit slowly depolarizing membrane potential ramps (~10 s) preceding the onset of spontaneous movement in both superficial and deep neurons. In spiking neurons, membrane potential ramps are accompanied by slow firing rate ramps with similar dynamics. In animals trained in a goal-directed go/no-go task in a virtual-reality environment, these membrane potential and spike rate ramps are accelerated during movement preparation. To assess the role of different interneuron subpopulations in preparatory activity in MOs, we chemogenetically suppressed the activity of local PV+ or SOM+ cells in MOs while recording from principal neurons during spontaneous movement periods, unveiling distinct roles for different subtypes of interneurons in shaping task-dependent membrane potential and firing rate ramps preceding the onset of movement.

## RESULTS

To explore neuronal dynamics preceding movement onset across different behavioral tasks, we used a setup adapted for rodent head-fixed navigation. Two groups of mice were subject to different behavioral paradigms: a control group of animals performed self-paced spontaneous movement on a treadmill in a dark environment after a brief habituation period, while another group of animals was trained in a goal-directed behavioral task in a virtual-reality (VR) environment (Figure 1A, see Methods). This latter group of animals learned to stop in a reward zone at the end of a linear VR corridor within ~6 days of training, as quantified for example by increased reward and success rates (Figure 1B-C). To establish the role of MOs in this goal-directed task, we inactivated MOs by local infusion of muscimol (Figure 1F-H, see Methods), and found that muscimol application significantly and reversibly reduced behavioral performance, such as reward and success rates (Figure 1D-E and 1G-H), without substantially affecting overall movement patterns of the animal (Figure 1E), confirming a specific role for MOs in the goal-directed behavioral task (Barthas and Kwan, 2017; Coen et al., 2021).

**Figure 1.**
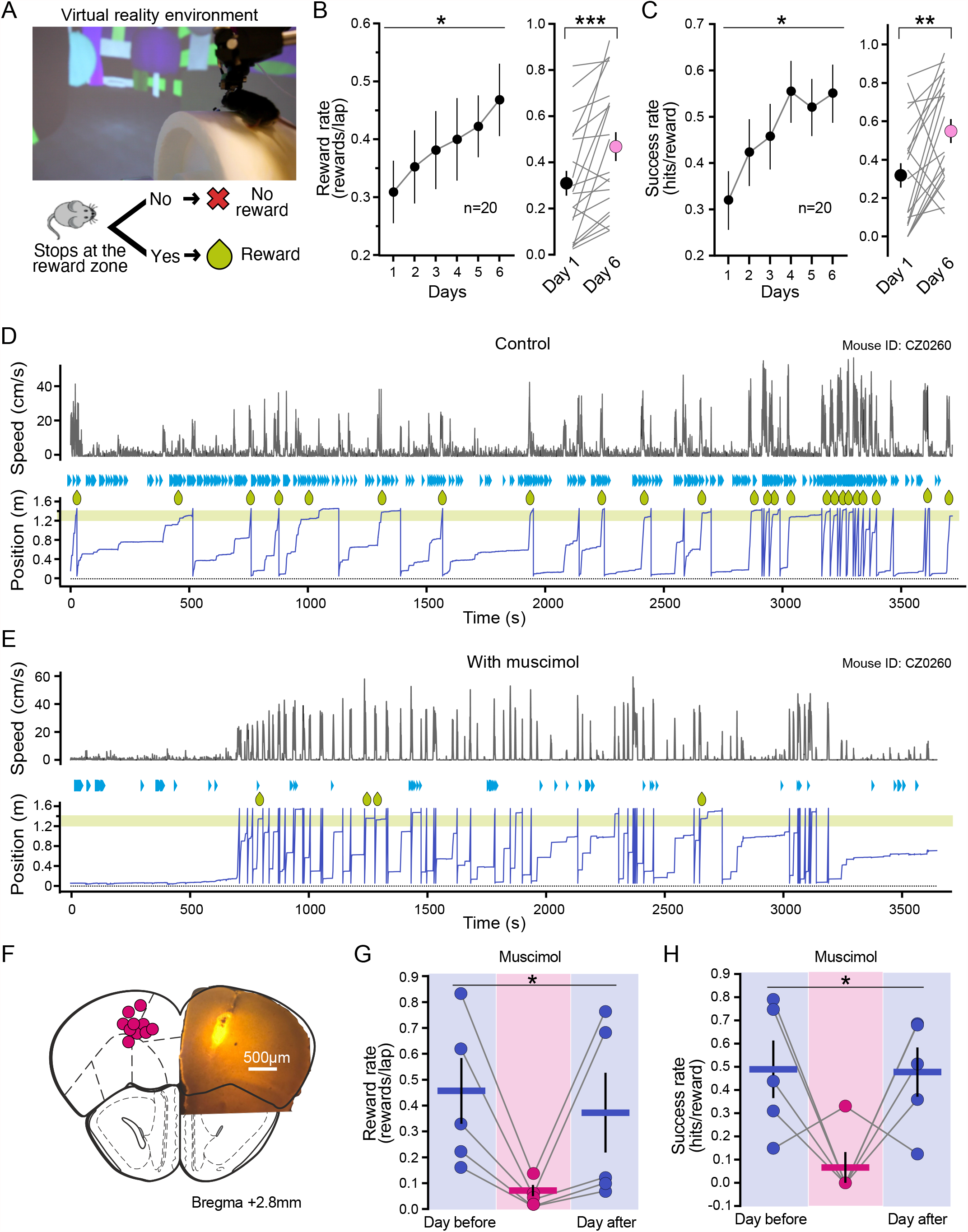
Role of MOs in a goal-directed behavioral VR task. (A) Top, The image shows a head-fixed mouse running in a virtual-reality (VR) environment. Bottom, Schematic drawing of the goal-directed go/no-go task. (B-C) Training performance quantified as reward rate (B, dispensed rewards per lap) and success rate (C, successful licks (“hits”) per dispensed reward). In each panel, the left graph shows training performance across days, the right graph compares training performance on day 1 vs day 6 (reward rate: day 1, 0.31 ± 0.04 vs day 6, 0.47 ± 0.05 rewards/lap; success rate: day 1, 0.31 ± 0.05 vs day 6, 0.55 ± 0.05 hits/reward; *n* = 20 mice). (D-E) Example training sessions from the same mouse under control conditions (D) and after muscimol application the next day (E). Muscimol was applied locally to MOs through a cannula 1 hour before the task. Top traces show animal speed, bottom traces show animal position on the virtual-reality track, green drops indicate dispensed rewards, blue triangles indicate licks. The reward zone is located between 1.2–1.4 m along the track (green shaded region). (F) Cannula tip positions (left) and coronal section (right) showing bodipy fluorescence in MOs where muscimol was injected bilaterally (0.6 µg/µL, 350 nL per site). (G-H) MOs inactivation by muscimol reversibly disrupts task performance, as quantified by the reward rate (G) and the success rate (H) (reward rate: day before 0.46 ± 0.14 vs muscimol 0.07 ± 0.02 vs day after 0.37 ± 0.17 rewards/lap; success rate: day before 0.49 ± 0.14 vs muscimol 0.07 ± 0.07 vs day after 0.48 ± 0.12 hits/reward; *n* =5 mice). Error bars represent s.e.m. Statistical significance was assessed using repeated measures ANOVA (B, C, G, H), and Wilcoxon signed rank tests (B, C) for day 1 vs day 6. ns, not significant; \**p*<0.05; \*\**p*<0.01; \*\*\**p*<0.001. Statistic values are provided in Table S1.

To characterize intrinsic membrane properties of MOs principal neurons, we performed *in vivo* whole-cell patch-clamp recordings from head-fixed mice of the control group during resting states (Figure 2A-C). Neurons which met basic recording criteria (n=47, see Methods) were split into superficial (150-420 µm) and deep (430-850 µm) recordings according to their depth in MOs (Carlén, 2017; Franklin and Paxinos, 2019; Lein et al., 2007) (Figure 2B-D; see Methods). Consistent with previous *in vivo* recordings from other neocortical regions (Zhao et al., 2016), superficial MOs neurons (mean recording depth, 303 ± 16 µm; n = 24) differed significantly from deep neurons (547 ± 24 µm, n = 23) in intrinsic membrane properties, with deep neurons showing more depolarized baseline membrane potentials and higher excitability (Figure 2E-G, details in Table S1).

**Figure 2.**
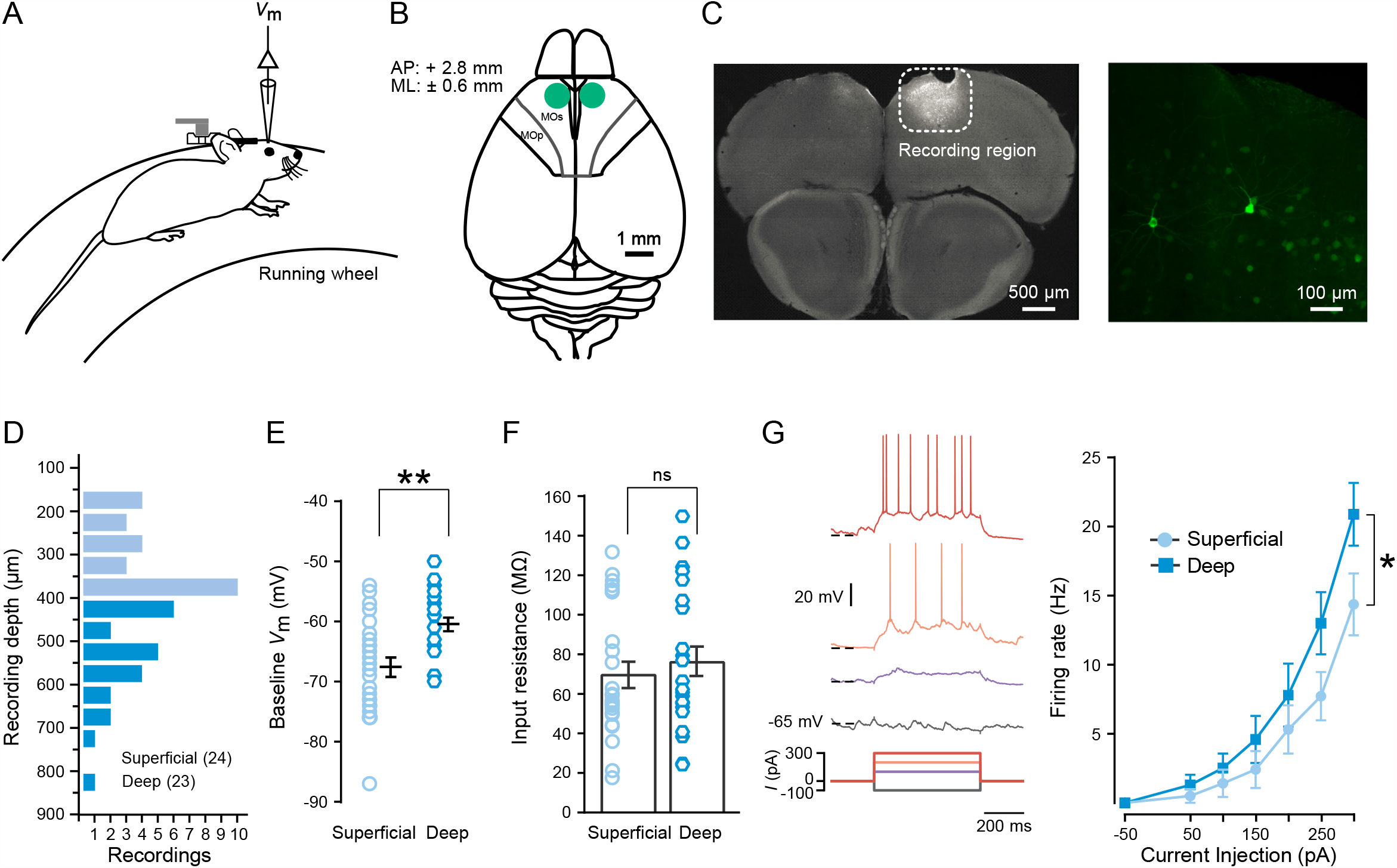
Distinct intrinsic membrane properties of superficial and deep MOs principal neurons *in vivo*. (A) Schematic drawing of the recording set-up. (B) Recording coordinates. (C) Left, coronal section of frontal cortex indicating the recording region in MOs (labeled by extracellular injection of tdTomato); right, biocytin-filled MOs principal neurons. (D) Distribution of recording depths (n=24 superficial neurons, n=23 deep neurons). (E) Deep MOs principal neurons show a more depolarized baseline membrane potential (*V*_m_) compared to superficial principal neurons. (F) No significant difference in input resistance between deep and superficial MOs principal neurons. (G) Left, Example *V*_m_ responses to sustained current injections. Right, relationship between firing rate and current injection (*f*-*I* curve). Deep neurons are more excitable than superficial neurons when large currents are injected (comparison between superficial and deep groups: *F*=9.94, *p*=0.002; comparision of two groups at 300 pA: *p*=0.01). Error bars represent s.e.m. Statistical significance was assessed using Mann-Whitney tests (D-F), and two-way ANOVA with Bonferroni post-hoc tests (G). ns, not significant; \**p*<0.05; \*\**p*<0.01. Mean values, s.e.m., and statistics details are provided in Table S1.

### Motion dependence of membrane potential and firing in MOs principal neurons

How does goal-directed behavior affect membrane potential and firing rate dynamics during resting and running states? To address this question, we recorded from MOs neurons in both groups of mice during locomotor behavior (Figure 3A). 29 out of the 47 recordings from the control group and all 18 of the recordings from the trained group reached the criteria for further analysis of motion-related membrane potential dynamics and firing patterns (see Methods). To compare motor behavior between the two groups, we quantified mean speed, time period *per* run and running frequency during the recordings. This analysis revealed that trained animals ran more frequently and at higher running speeds (Figure 3B–D). During spontaneous movement, we observed two populations of MOs neurons with distinct firing rate patterns during resting and running periods: most principal neurons (20 of 29 neurons, 70%) exhibited higher firing rates during resting periods, while a smaller group of neurons (7 of 29 neurons, 24%) showed higher firing rates during running periods (Figure 3G left: resting 0.50 ± 0.08 Hz *vs* running 0.54 ± 0.24 Hz, n = 29; 2 out of 29 neurons were not firing). Notably, in animals trained in the goal-directed task, we observed an opposite trend: more neurons (9 of 18 neurons, 50%) showed higher firing rates during running periods, while 8 of 18 neurons (44%) showed lower firing rates during running than resting periods (Figure 3G right: resting 1.21 ± 0.33 Hz *vs* running 2.42 ± 0.80 Hz, n = 18; 1 out of 18 neurons was not firing). Detailed analysis of membrane potential dynamics revealed that mean membrane potential was more depolarized during running than resting periods in both groups of animals (Figure 3H). The membrane potential increase between resting and running periods was significantly larger during the goal-directed task (Figure 3I). Across neurons, we did not observe any consistent correlation between membrane potential change and animal speed, indicating that state-dependent membrane potential changes cannot be explained by a simple linear relationship with animal speed (Figure S1).

**Figure 3.**
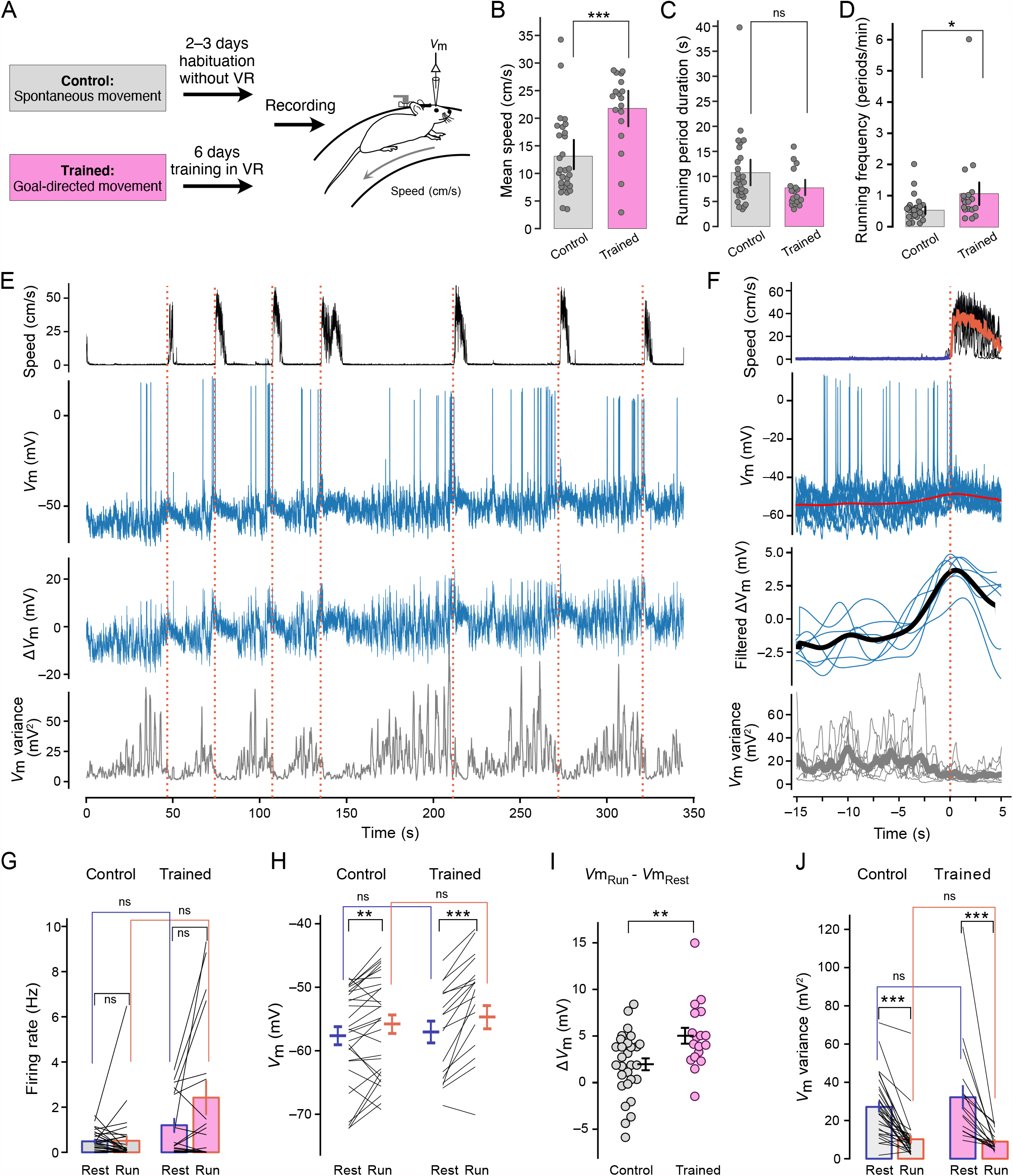
Differences in subthreshold membrane potential and firing rates between resting and running periods in MOs neurons. (A) Experimental timeline for spontaneously running control mice (top) and mice trained in the goal-directed task (bottom). (B-D) Comparison of the mean running speeds (B), duration of individual running periods (C), and rate of running periods (D) between recordings from the control and trained groups (mean speed: control 13.1 ± 1.4 vs trained 21.8 ± 1.7 cm/s; running period duration 10.8 ± 1.4 vs 7.7 ± 0.9 s; running frequency: control 0.6 ± 0.1 vs trained 1.1 ± 0.3 periods per minute; *n* = 29 for control and *n* = 18 for trained groups). Symbols represent individual recordings. (E) Example whole-cell recording from a superficial MOs neuron during goal-directed behavior. Traces show (from top) animal speed, membrane potential (*V*m), *V*m after blanking action potentials, and *V*m variance. Vertical dashed lines indicate movement onset. (F) Same recording as in (E), with data aligned to the onset of running periods at *t* = 0s. Traces show (from top) animal speed, membrane potential (*V*m), low-pass filtered *V*m after blanking action potentials, and *V*m variance. Thin traces represent individual running periods, thick traces represent the mean across running periods. The thick trace on top of *V*m (second from top) represents the mean of low-pass filtered *V*m after blanking action potentials. (G–J) Summary of firing rates (G), mean *V*m (H), membrane potential difference (Δ*V*_m_) between running and resting periods (I), and *V*m variance (J) during resting (blue) and running periods (red), for recordings from the control group (*n* = 29) and from the group of mice trained in a goal-directed task (*n* = 18) (H, left: control group: resting −58.0 ± 1.5 mV vs running −56.0 ± 1.5 mV; H, right: trained group: resting −57.0 ± 1.7 mV vs running −51.9 ± 1.8 mV; I: control group: 1.98 ± 0.65 mV; trained group: 5.04 ± 0.87 mV; J, left: resting 26.5 ± 3.0 mV^2^ vs running 9.5 ± 2.2 mV^2^; right: resting 32.2 ± 6.5 mV^2^ vs running 9.0 ± 1.2 mV^2^). Error bars represent ± s.e.m. Statistical significance was assessed using Wilcoxon signed rank tests (G, H and J) for paired groups, and Mann-Whitney tests (B-D, and G-J) for unpaired groups. ns, not significant; \**p*<0.05; \*\**p*<0.01; \*\*\**p*<0.001. Mean values, s.e.m., and statistics details are provided in Table S1.

Several previous studies have revealed that the amplitude of membrane potential fluctuations in neocortical neurons decreases promptly upon changes of behavioral states, reflecting a rapid transition from a synchronized to a desynchronized cortical state (Bennett et al., 2013; Churchland et al., 2010a; Eggermann et al., 2014; Polack et al., 2013; Poulet and Petersen, 2008; Poulet et al., 2012; Schiemann et al., 2015; Schneider et al., 2014; Zagha et al., 2013; Zhou et al., 2014). In agreement with these findings, we found that MOs neurons displayed larger subthreshold membrane potential fluctuations during resting periods compared to running periods in the control group (Figure 3J). This decrease in membrane potential fluctuation amplitude was also observed during goal-directed running, similar to the untrained group of animals (Figure 3E-F and 3J). These fluctuations contained broadband frequency components without any obvious peak in the spectrum before and during movement (Figure S2). Thus, our data indicate that membrane potential fluctuations in MOs neurons rapidly transition from large to small amplitudes, reflecting a change to a desynchronized low-variability state upon movement onset (Churchland et al., 2010a) (Figure 3E-F, 3J and S2).

### Temporal dynamics of preparatory ramping signals preceding the onset of movement

To characterize the membrane potential dynamics underlying pre-movement spiking activity, we analyzed recordings with sufficiently long recording periods before and after onset of movement from neurons spontaneously spiking preceding running periods (Figure 4; *n* = 11 out of 29 recordings matching criteria; see Methods). Changes in subthreshold membrane potential (Δ*V*_m_) and firing rates were aligned to the onset of spontaneous running periods of untrained animals (Figure 4A, C). This analysis revealed that during spontaneous movement, subthreshold membrane potential displayed a gradual depolarization (~10 s) preceding running onset (Figure 4A, individual examples; Figure 4C, summary data). Simultaneously, firing rates averaged across different animals showed slowly and gradually increasing firing rates during preparation of movement (Figure 4C). To further probe how these depolarizing membrane potential ramps affect firing patterns in larger populations of neurons during preparation of movement, we performed extracellular recordings of MOs population activity with Neuropixels probe from one untrained mouse (Figure S3A, B). These recordings revealed firing rate ramps in putative principal neurons preceding spontaneous movement with similar temporal dynamics as observed in our whole-cell recordings (Figure S3C, D). In particular, a steady increase in pre-motion firing rates could be observed in sparsely firing neurons (Figure S3E) that matched the mean firing rates observed during whole-cell recordings (Figure 4C).

**Figure 4.**
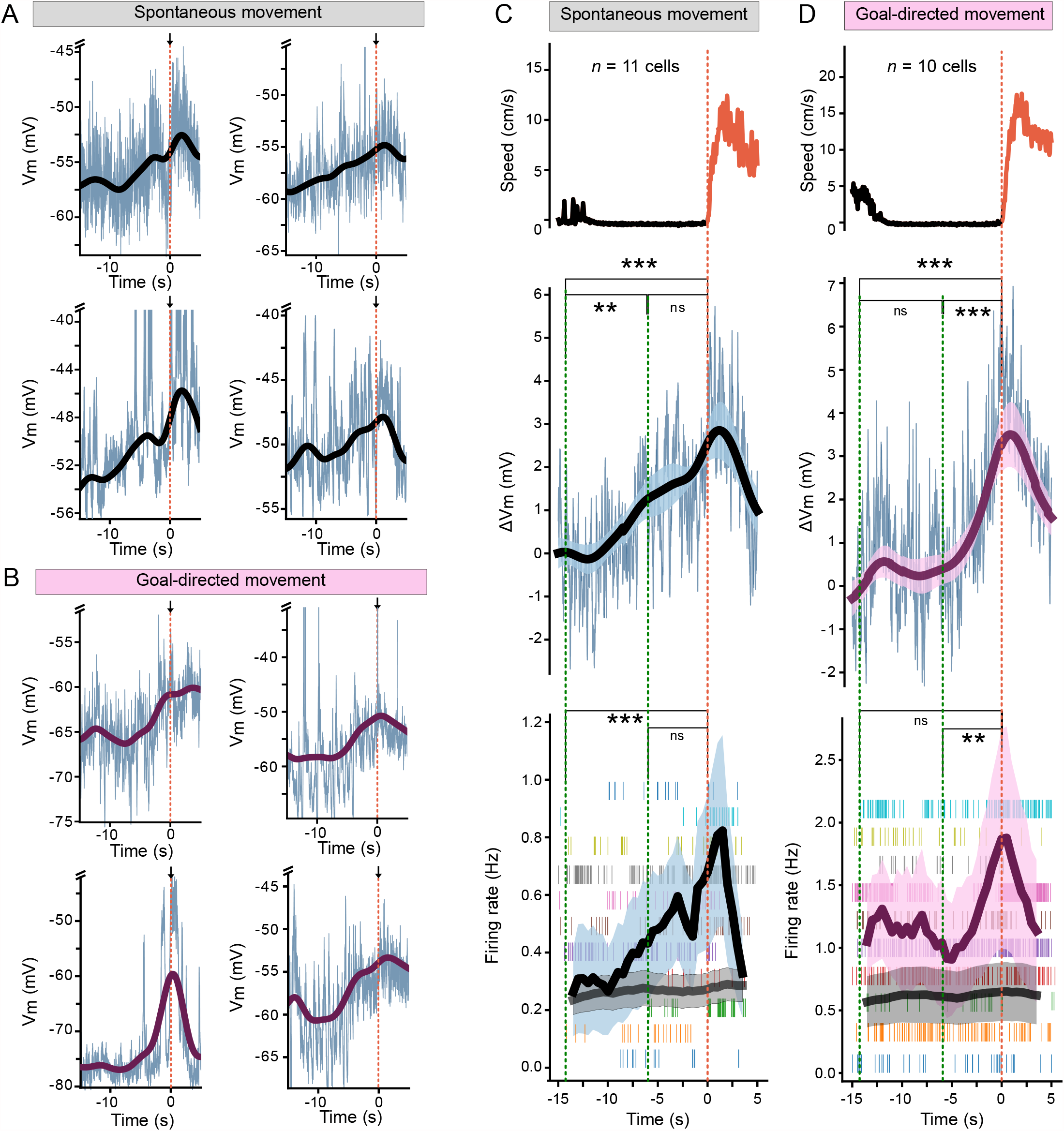
Task dependence of membrane potential and firing rate dynamics preceding movement onset. (A-B) Example whole-cell recordings from different MOs neurons preceding movement onset at *t* =0s (indicated bya red vertical dashed line and an arrow). Thin blue traces represent raw membrane potential, thick traces represent low-pass filtered membrane potential after blanking action potentials. Note slower dynamics of depolarizing ramps in control animals (A) compared to trained animals (B). (C-D) Summary of speed, membrane potential and firing rate dynamics preceding movement onset across control animals (C, n=11 recordings) and trained animals (D, n=10 recordings). Data are aligned to running onset at t = 0 s (indicated by a red vertical dashed line). Top, mean animal speed. Middle, mean membrane potential (thin traces), mean low-pass filtered membrane potential (thick traces, shaded regions represent mean ± s.e.m). Bottom, thick traces represent mean spike firing rates (black), and mean of shuffled firing rates (grey, n=100 shuffles). Shaded regions represent mean ± s.e.m. Ticks represent spike firing, with all running periods of a single recording shown in a single row. Statistical significance was assessed by Spearman’s correlation (C-D). ns, not significant; *p<0.05; **p<0.01; ***p<0.001. Statistics details are provided in Table S1.

To probe how goal-directed training affects synaptic integration during movement preparation, we also analyzed membrane potential and firing rates preceding movement in whole-cell recordings from animals trained in the goal-directed task (*n* = 10 out of 18 recordings matching criteria; see Methods). Strikingly, after training, membrane potential ramps were accelerated (~6 s) (Figure 4B, individual examples; Figure 4D, summary data). Simultaneously, we observed faster spike ramps with a larger amplitude during preparation of movement in these recordings (Figure 4D). Thus, the dynamics of both sub- and suprathreshold ramping activity appears to depend on the nature of the behavioral task.

To identify possible behavioral correlates of the depolarizing membrane potential ramps, we captured body and whisker movements of animals across several sessions during goal-directed training in the VR environment (Figure S4A). Inspection of the training videos showed that whisker movement often preceded running periods (Figure S4B; see Methods). Detailed quantification of these whisker precession times revealed that they depended on the training state of the animal, with longer whisker precession times in untrained compared to trained animals (Figure S4C). Across ~6 days of training sessions, whisker precession times decreased from ~9s to ~6 s (Figure S4D), comparable to the temporal dynamics of membrane potential and firing rate ramps preceding running onset (Figure 4).

Previous work has revealed differences in preparatory neuronal activity between neurons in superficial and deep layers of the frontal motor cortex in a whisker-based task (Chen et al., 2017; Wagner et al., 2019). As recordings from superficial and deep neurons in our data set showed differences in baseline membrane potential and intrinsic membrane properties (Figure 2E-G), we analysed superficial and deep recordings separately (Figure S5-S6). This analysis revealed more pronounced changes in membrane potential and firing rates after behavioral training in deep compared to superficial neurons (Figure S6A–C), while pre-movement membrane potential dynamics were comparable across layers (Figure S6D–I).

### Local inhibitory neurons disinhibit principal neurons and shape membrane potential ramps

Which cellular and circuit processes could mediate the sustained depolarization of membrane potential in MOs neurons during preparation of movement? Emerging evidence suggests that disinhibition plays a critical role in enhancing neuronal activity during diverse behavioral functions (Letzkus et al., 2015; Wolff et al., 2014). To test the role of disinhibition of MOs neurons, we chemogenetically suppressed the activity of local PV+ or SOM+ interneurons in MOs (see Methods) while recording from principal neurons during spontaneous movement periods (Figure 5A-B). In line with previous reports (Jackson et al., 2018), chemogenetic inactivation of PV+, but not of SOM+ interneurons, resulted in an increase of basal firing rates (Figure 5D) and membrane potential fluctuations (Figure 5F) of MOs principal neurons, without affecting the mean membrane potential (Figure 5E) when animals were resting on the treadmill. Chemogenetic inactivation of PV+, but not SOM+ interneurons, increased the movement-related firing rates of MOs principal neurons during running periods (Figure 5G). When we inactivated PV+ interneurons, we observed that 9 out of 14 principal neurons (64%) exhibited higher firing rates during running compared to resting periods (6 out of 9 superficial recordings, and 3 out of 5 deep recordings; details in table 1). During inactivation of SOM+ interneurons, 6 out of 12 principal neurons (50%) exhibited higher firing rates during running periods (6 out of 9 superficial recordings, and 0 out of 3 deep recordings). However, a larger depolarization of principal neurons during transition between resting and running periods was only observed after inactivation of PV+, but not of SOM+ interneurons (Figure 5H-I). By contrast, we still observed a decrease of membrane potential fluctuations when either PV+ or SOM+ interneurons were inactivated (Figure 5J). Thus, our results suggest that PV+ and SOM+ interneurons produce differential effects on membrane potential and firing rates of principal neurons during different locomotor states.

**Figure 5.**
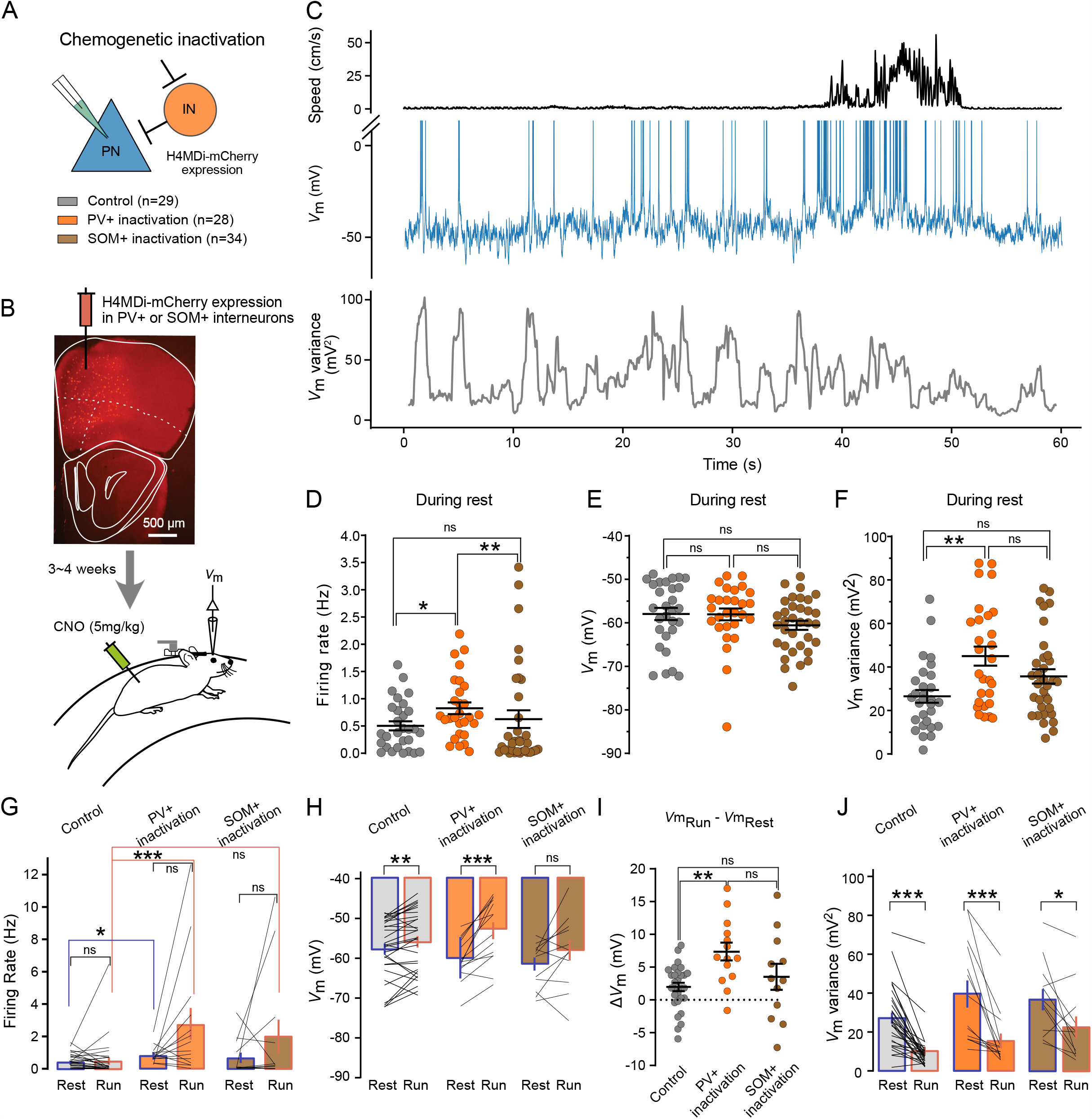
Inactivation of local PV+ interneurons, but not of SOM+ INs, in MOs affects movement-related firing patterns. (A) Schematic drawing of the experimental paradigm. Control recordings are the same as those shown in Figure 3. (B) Fluorescence image of h4MDi-mCherry-expressing SOM+ interneurons in a coronal slice of MOs. CNO was injected i.p. 1h before recordings to activate H4MDi receptors. Experiments were performed within 3h after CNO injection. (C) Example intracellular recording from a MOs neuron during chemogenetic inactivation of PV+ interneurons. Traces show (from top) animal speed, membrane potential (*V*_m_), and *V*_m_ variance. (D–F) Comparison of firing rates (D), mean membrane potential (E), and *V*_m_ variance (F) under control conditions (n=29) and during chemogenetic inactivation of PV+ (*n* = 28) or SOM+ (*n* = 34) interneurons during resting periods (D: control conditions represent the same data as shown in Figure 3G, left, rest; during PV+ inactivation: 0.82 ± 0.11 Hz; during SOM+ inactivation: 0.63 ± 0.17 Hz; E: control conditions represent the same data as shown in Figure 3H, left, rest; during PV+ inactivation: −58.1 ± 1.3 mV; during SOM+ inactivation: −60.6 ± 1.1 mV; F: control conditions represent the same data as shown in Figure 3J, left, rest; during PV+ inactivation: 45.1 ± 4.3 mV^2^; during SOM+ inactivation: 35.6 ± 3.6 mV^2^). (G–J) Comparison of firing rates (G), membrane potential (H), changes in membrane potential (I), and *V*_m_ variance (J) during resting (blue) and running periods (red) under control conditions (*n* = 29) and during chemogenetic inactivation of PV+ (*n* = 14), or SOM+ interneurons (*n* = 12). Recordings in G–J are a subset of the recordings in D–F that include running periods. (G: control conditions represent the same data as in Figure 3G, left; during PV+ inactivation: resting 0.86 ± 0.14 Hz vs running 2.77 ± 1.02 Hz; during SOM+ inactivation: resting 0.71 ± 0.33 Hz vs running 2.07 ± 1.10 Hz; H: control conditions represent the same data as shown in Figure 3H; PV+ inactivation: resting −60.3 ± 1.4 mV vs running −53.0 ± 2.2 mV; SOM+ inactivation: resting −61.8 ± 1.7 mV vs running −58.3 ± 2.7 mV; J: control conditions represent the same data as shown in Figure 3J, left; PV+ inactivation: resting 39.9 ± 7.2 mV^2^ vs running 15.9 ± 4.0 mV^2^; SOM+ inactivation: resting 37.2 ± 5.7 mV^2^ vs running 23.1 ± 5.8 mV^2^). Error bars represent ±s.e.m. Statistical significance was assessed using Kruskal-Wallis tests (D-F and I), Wilcoxon signed rank tests (H, I and K) for paired groups, and Mann-Whitney tests (G) for unpaired groups. ns, not significant; \**p*<0.05; \*\**p*<0.01; \*\*\**p*<0.001. Mean values, s.e.m., and statistics details are provided in Table S1.

We next sought to assess the differential roles of SOM+ and PV+ cells in shaping preparatory membrane potential ramp dynamics. To consistently compare the effects of inactivation of different interneurons with our control data, we focused only on recordings with sufficiently long recording durations before and after running periods from principal neurons spontaneously spiking preceding running periods (Figure 6; n = 8 spiking recordings during inactivation of PV+, depth: 338 ± 48 µm; n = 7 spiking recordings during inactivation of SOM+, depth: 359 ± 41 µm). We observed that during inactivation of PV+ interneurons, both subthreshold membrane potential and firing rate ramps were still preserved, without substantial changes in their duration (~10 s) (Figure 6A, C). By contrast, membrane potential and firing rate ramps were abolished by inactivation of SOM+ interneurons (Figure 6B, D). When we compared membrane potential ramps between inactivation of interneurons and control recordings, we found that inactivation of PV+ cells increased the amplitude of ramps (Figure 6C and S7A), while inactivation of SOM+ interneurons depressed the ramps (Figure 6D and S7B). Notably, during preparation of movement, we observed a sustained depolarization of membrane potential in MOs principal neurons during inactivation of PV+, but not of SOM+ interneurons, compared to control recordings (Figure S7C-D). Thus, our data show that inactivation of PV+ interneurons leads to depolarized, large-amplitude membrane potential ramps during spontaneous running periods, resembling those observed after goal-directed training (Figure 4C-D, 6C and S7C). Moreover, inactivation of SOM+ cells abolishes slow membrane potential ramps (Figure 6D) without changing the baseline membrane potential (Figure S7D). In addition, extracellular recordings from putative interneurons revealed specific changes in their firing activity both before and after the onset of spontaneous running periods (Figure S3F-G). While these recordings do not distinguish between PV+ and SOM+ interneurons, they generally support the view that interneurons play specific roles during movement preparation.

**Figure 6.**
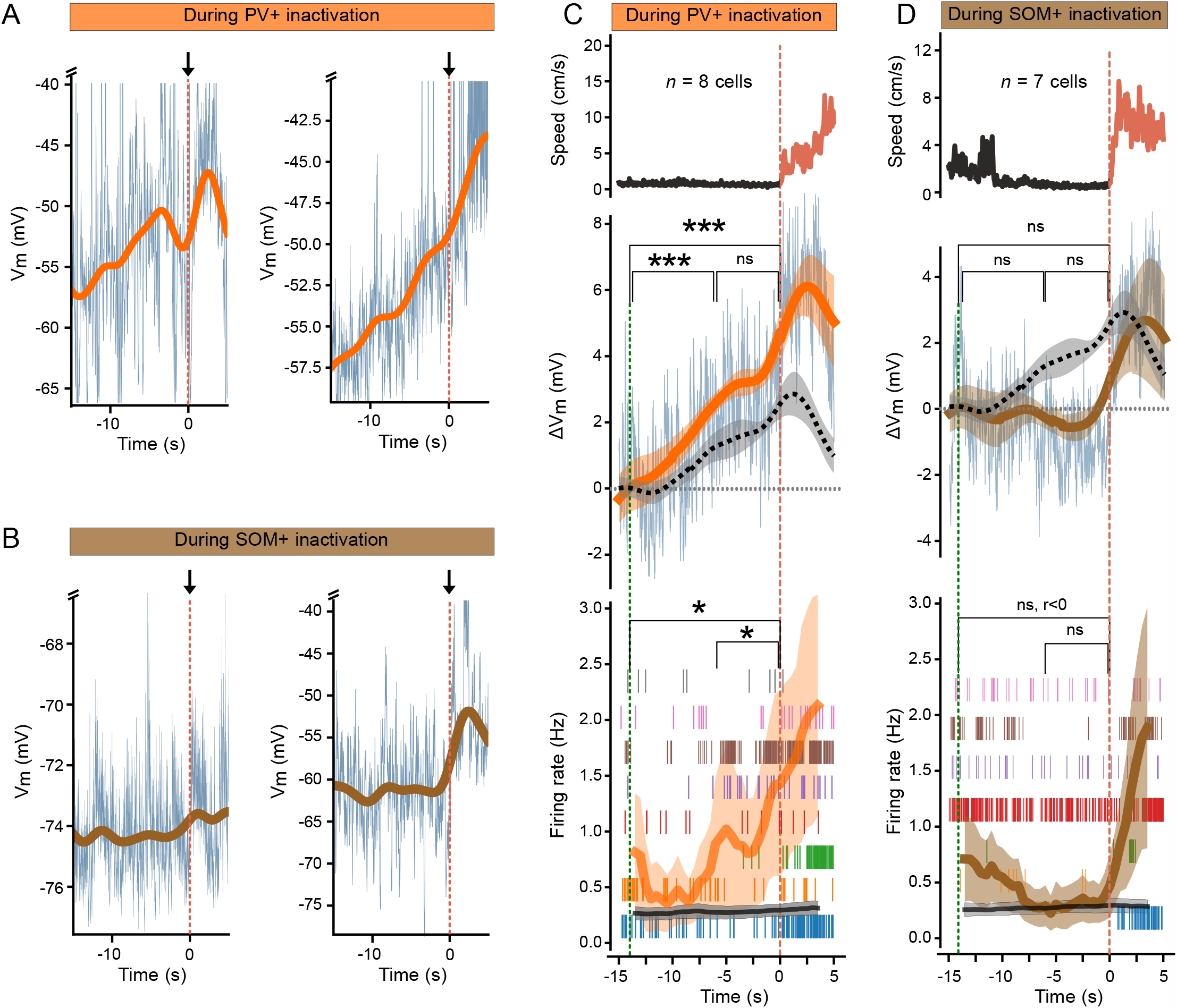
Activity of local PV+ and SOM+ interneurons in MOs shapes subthreshold membrane potential ramps. (A-B) Example whole-cell recordings from different MOs neurons preceding movement onset at t = 0 s (indicated by a red vertical dashed line and an arrow) during chemogenetic inactivation of PV+ interneurons (A) or of SOM+ interneurons (B). Thin blue traces represent raw membrane potential, thick traces represent low-pass filtered membrane potential. (C-D) Summary of speed, membrane potential and firing rate dynamics preceding movement onset during chemogenetic inactivation of PV+ interneurons (C, n=8 recordings) or of SOM+ interneurons (D, n=7 recordings). Data are aligned to running onset at t = 0 s (indicated by a red vertical dashed line). Top, mean animal speed. Middle, mean membrane potential (thin traces), mean low-pass filtered membrane potential (thick traces, shaded regions represent mean ± s.e.m). The black dash traces represent the control recordings (n=11 spiking neurons, same as shown in Figure 4C). Bottom, thick traces represent mean spike firing rates (orange and brown), and mean of shuffled firing rates (grey, n=100 shuffles). Shaded regions represent mean ± s.e.m. Ticks represent spike firing, with each row showing a different recording. Statistical significance was assessed by Spearman’s correlation. ns, not significant; *p<0.05; **p<0.01; ***p<0.001. Statistics details are provided in Table S1.

### Concerted action of external inputs and local inhibition drives task-dependent preparatory motor signals

In cortical circuits, PV+ interneurons mainly exert fast and powerful perisomatic inhibition on principal neurons and other PV+ cells, while SOM+ interneurons mainly form inhibitory synapses onto distal dendrites of principal neurons and PV+ cells (Cottam et al., 2013; Hangya et al., 2014; Hu et al., 2014; Pfeffer et al., 2013; Rudy et al., 2011). To obtain a better understanding of the differential role of SOM+ and PV+ interneurons in shaping preparatory neuronal dynamics, we developed a simple model of the local MOs circuit (Figure 7; see details in Table S2). PV+, SOM+ and principal neurons were modeled as rate-based variables. All 3 types of neurons received external thalamocortical input (Ährlund-Richter et al., 2019) that increased in activity in a step-like manner during movement preparation. The SOM+ cell inhibited both the PV+ interneuron and the principal neuron, whereas the PV+ cell mainly inhibited the principal neuron. The principal neuron excited both interneuron units and itself in a recurrent manner. As has previously been shown (Chaudhuri and Fiete, 2016; Economo et al., 2018; Gao et al., 2018; Li et al., 2016; Murakami et al., 2014), the recurrent connectivity of the principal neuron, in concert with inhibition from PV+ and SOM+ cells, resulted in a conversion of the step-like external input into a slowly increasing and sustained ramp of activity (Figure 7A), which is consistent with our experimental observations (Figure 4 and S6). Inactivation of the SOM+ cell led to disinhibition of the PV+ interneuron and consequently to silencing of the principal neuron, thereby abolishing the ramp (Figure 7B). By contrast, inactivation of the PV+ cell led to disinhibition of the principal neuron, resulting in a ramp with a steeper slope and in increased baseline activity (Figure 7C). Both of these simulation results were qualitatively consistent with our experimental observations of membrane potential dynamics during inactivation of PV+ or SOM+ interneurons (Figure 6 and S7). Our simulations further revealed that increasing excitatory synaptic weights on the SOM+ cell could accelerate the temporal dynamics and amplitude of the ramping activity by changing the SOM+/PV+ activity balance over time (Figure 7D). These simulations show that the interplay between SOM+ and PV+ interneuron activities can explain their roles in shaping the ramping dynamics in principal neurons, in agreement with our experimental findings (Figure 6 and S7). Moreover, our model reveals that potentiation of excitatory synapses onto SOM+ cells changes the balance between SOM+ and PV+ interneuron activities during preparation of movement, leading to decreased PV+ activity during movement preparation and during running (Figure 7D). These dynamics can thereby contribute to the acceleration of the ramp that we observe after goal-directed training (Figure 4 and S6).

**Figure 7.**
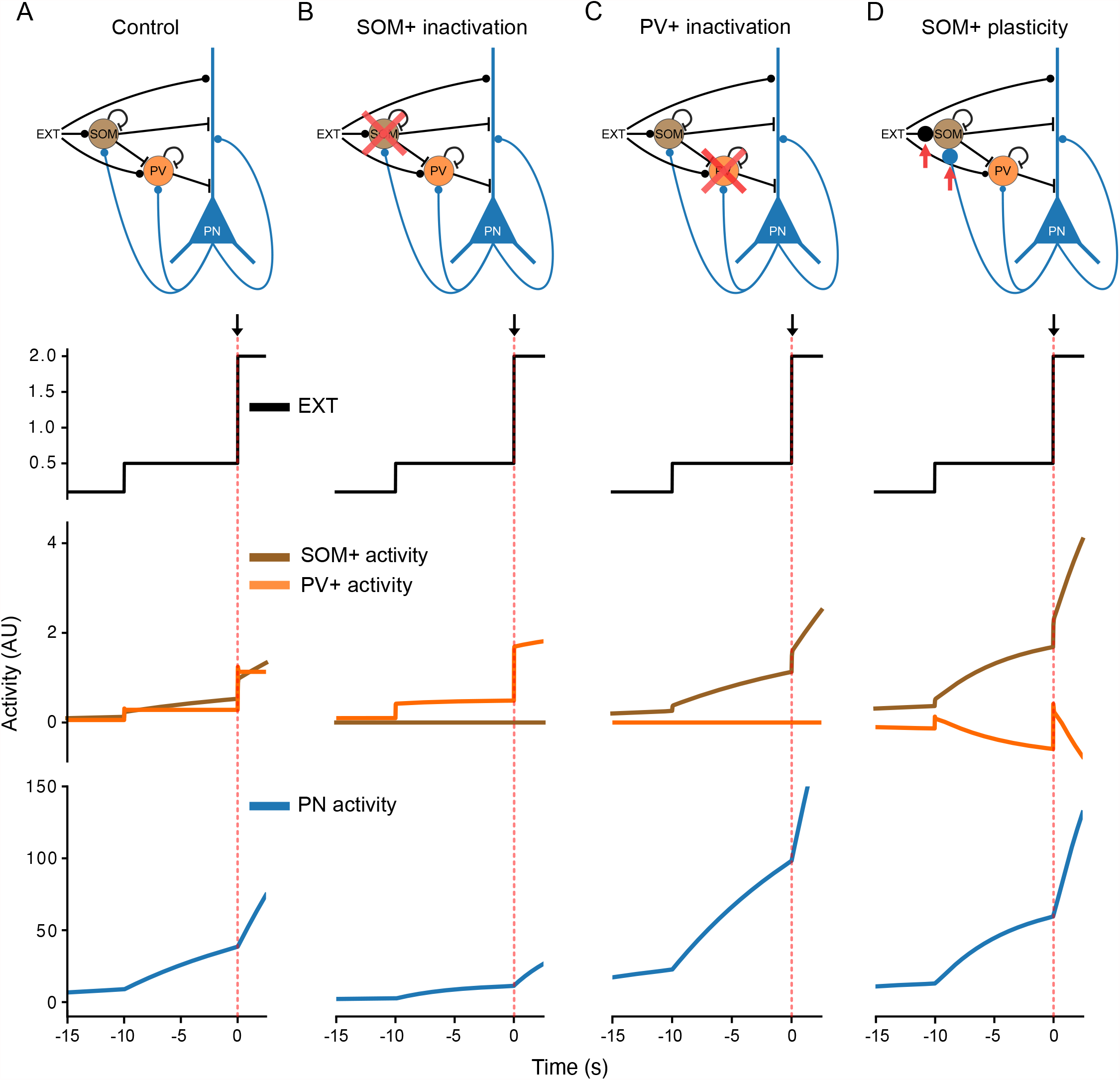
Differential role of SOM+ and PV+ interneurons during movement anticipation in a computational model of the local MOs circuit. (A) Top, A simple computational model of the local MOs circuit with 3 units: SOM+ interneurons, PV+ interneurons, and principal neurons (PNs). All units receive the same external inputs. Bottom, simulation results. Activity in arbitrary units is plotted against time. External inputs (top) increase in a step-like fashion during movement anticipation at *t* = –10 s. By virtue of their recurrent connectivity in conjunction with inhibitory inputs (middle), pyramidal cells (bottom) respond with a graded slow increase of activity. (B) Simulation of chemogenetic inactivation of SOM+ interneurons. Absence of SOM+-mediated inhibition of PV+ interneurons leads to an increase in PV+ activity, thereby abolishing the ramp in pyramidal neurons. (C) Simulation of chemogenetic inactivation of PV+ interneurons. Absence of inhibition from PV+ interneurons leads to an acceleration of ramping activity, and to an increase of baseline activity. (D) Simulation of increasing excitatory synaptic weights on SOM+ interneurons. Increased activation of SOM+ interneurons leads to a change in SOM+/PV+ activity balance over time in favor of SOM+ interneurons, thereby accelerating the ramping activity in principal neurons.

## DISCUSSION

Using *in vivo* whole-cell patch-clamp recordings from principal neurons in the secondary motor cortex, we explore how excitatory and inhibitory synaptic inputs are integrated during preparation of motor behavior. We observe that both superficial and deep neurons display different neuronal activity dynamics depending on whether the animal was trained to perform a behavioral task: untrained animals show slow (~10 s) ramps of membrane potential and spike rates preceding spontaneous movement periods, whereas in animals trained to perform a goal-directed task, the dynamics of both membrane potential and spike ramps are faster (~6 s) and larger in amplitude. At the same time, membrane potential fluctuations rapidly decrease in amplitude upon onset of running, independently of the training state of the animal. To understand how these dynamics are generated at the cellular and circuit level, we manipulated the activity of specific interneuron subpopulations using chemogenetic tools. Inactivation of PV+ interneurons disinhibits MOs principal neurons and increases the amplitude of membrane potential ramps, while inactivation of SOM+ cells abolishes membrane potential ramps. However, local inactivation of PV+ or SOM+ interneurons does not affect the running-related decrease in membrane potential fluctuation amplitude. Therefore, our results suggest that the concerted action of external inputs and local inhibition shapes preparatory motor signals in MOs.

### Circuit mechanisms underlying subthreshold membrane potential fluctuations before and after onset of movement

Several mammalian brain regions transition between synchronized and desynchronized regimes when adapting to changes between different behavioral states or responding to different stimuli (Buzsáki and Draguhn, 2004; Churchland et al., 2010a; Harris and Thiele, 2011; Lee and Dan, 2012). Slow subthreshold fluctuations during synchronized states have previously been observed during resting periods in somatosensory, visual, and auditory cortices of head-restrained mice. When animals start to move or attend to a stimulus, these cortical neurons rapidly transition to a desynchronized low-variability state (Bennett et al., 2013; Churchland et al., 2010a; Eggermann et al., 2014; Polack et al., 2013; Poulet and Petersen, 2008; Poulet et al., 2012; Schiemann et al., 2015; Schneider et al., 2014; Zagha et al., 2013; Zhou et al., 2014). Consistently, we observed that most MOs principal neurons showed large subthreshold fluctuations during resting periods, and transitioned to a low-fluctuation state during running periods (Figure 3J and S6C). Thus, our results suggest that neurons in MOs evolve from a synchronized to a desynchronized state upon running onset.

Which mechanisms can explain membrane potential fluctuation variability? Membrane potential fluctuations are governed by the interplay between intrinsic membrane properties and excitatory and inhibitory synaptic inputs. Excitatory and inhibitory neurons in the barrel cortex have been shown to affect subthreshold fluctuations during quiet wakefulness (Gentet et al., 2010). In our experiments, chemogenetic inactivation of PV+, but not of SOM+ interneurons, results in higher firing rates and increased subthreshold membrane potential fluctuations during resting periods (Figure 5D-F). Local PV+ cells in the auditory cortex have been suggested to play a role in controlling motion-related membrane potential fluctuation amplitudes (Schneider et al., 2014). However, we observed that chemogenetic suppression of local PV+ interneurons reversed the motion-related firing pattern of principal neurons, but left the decrease of subthreshold fluctuations upon movement onset unaffected (Figure 5J). Thus, our results support the view that the motion-related decrease in membrane potential fluctuations is driven by a desynchronization of external inputs (Churchland et al., 2010a) rather than by increased activity of local interneurons.

What could be the source of these external inputs? It has been suggested that coordinated activity in a multiregional loop spanning cerebellum, thalamus and frontal cortex is required for anticipatory activity (Gao et al., 2018; Guo et al., 2014, 2017; Wagner et al., 2019). Activity in the cerebellum is necessary to maintain preparatory activity in the frontal cortex, which in turn affects the preparatory activity in deep cerebellar nuclei (Chabrol et al., 2019; Gao et al., 2018). At the anatomical level, frontal cortex projects to the cerebellum via the basal pontine nucleus, and cerebellum projects back to frontal cortex via the thalamus (Ährlund-Richter et al., 2019; Economo et al., 2018; Gao et al., 2018; Guo et al., 2017; Li et al., 2015). Therefore, thalamic projections to the neocortex play a crucial role in driving cortical sensory processing (Fuster and Alexander, 1971; Lara et al., 2018). How do these thalamo-cortical synapses modulate movement-related membrane potential dynamics? In a goal-directed motor task, silencing thalamic inputs to primary motor cortex blocks characteristic movement-related membrane potential dynamics and movement initiation (Dacre et al., 2021). Similarly, during whisking, membrane potential depolarization and desynchronization of the cortical state are driven by increased thalamic activity (Dacre et al., 2021; Poulet et al., 2012). These previous studies suggest that thalamic inputs may also drive the preparatory activity that we observe in MOs as part of a feedback loop, and the concerted action of these external inputs with local inhibition shapes preparatory motor signals in MOs (Figure 7A).

### Information flow in the MOs circuit during preparation of movement

How individual neurons can generate preparatory signals during movement preparation is unclear (Churchland et al., 2010b; Inagaki et al., 2019). As previous recordings have focused on spiking neurons throughout the experiments, how synaptic inputs are integrated by MOs neurons during their silent periods in preparation of movement has not been revealed yet. Our intracellular recordings from silent and firing MOs neurons show that both types of cells integrate synaptic inputs to produce slowly depolarizing membrane potential ramps during preparation of movement (Figure 4 and S6). In cells that fire action potentials during preparation of movement, these membrane potential ramps translate into spike ramps with similar dynamics (Figure 4).

In the neocortex, information from the thalamus and cortical areas is transmitted in a directional manner from superficial to deep layers (Bureau et al., 2006; Lübke and Feldmeyer, 2007; Otsuka and Kawaguchi, 2008). Selective activation of deep layer 5 cells by superficial layer 2/3 neurons may facilitate this directional transfer of information (DeFelipe and Fariñas, 1992; Kampa et al., 2006; Otsuka and Kawaguchi, 2008). Superficial layers are thought to be the principal recipient for sensory information (Mao et al., 2011). By contrast, deep neurons in layer 5 of frontal areas produce movement output signals. For example, deep neurons from the anterior-lateral motor region play a distinct role in initiating ramping activity in preparation of movement (Chen et al., 2017; Guo et al., 2014). These movement signals are then sent along the thalamus and the corticospinal tract to the spinal cord and superior colliculus to drive movement output in coordination with the cerebellum (Donoghue and Wise, 1982; Economo et al., 2018; Gabbott et al., 2005). We find that both superficial and deep MOs neurons display slowly depolarizing membrane potential ramps preceding movement onset by several seconds (Figure S6D-I). However, deep neurons are more depolarized and excitable than superficial neurons (Figure 2D-G), rendering them more sensitive to changes in their inputs. Accordingly, neuronal spiking dynamics are more strongly affected by the training state of the animal in deep cells compared to superficial cells, as we observe a pronounced increase of firing rates only in deep neurons after behavioral tasks (Figure S6A). Together, our results support the view that excitatory synaptic inputs are processed in a feed-forward manner from superficial to deep layers, resulting in task-dependent output from deep cortical neurons that exerts top-down influences in information processing (Gilbert and Sigman, 2007).

### Inhibitory role of PV+ interneurons during preparation of movement

After goal-directed training, MOs neurons were more depolarized throughout preparation of movement compared to neurons from spontaneously running animals (Figure S6F, I). At the same time, the mean membrane potential during resting periods was not significantly different between the two groups (Figure 3H). How can we explain the sustained seconds-long depolarization during movement preparation? Our chemogenetic experiments indicate that inactivation of PV+, but not of SOM+ interneurons, can result in a large and persistent depolarization of principal neurons during movement preparation and during running (Figure S7C), without changing the mean membrane potential during resting periods (Figure 5E). In addition, we also find that inactivation of PV+ cells can substantially increase membrane potential fluctuation amplitudes and firing rates during resting periods (Figure 5D, F). Furthermore, inactivation of PV+, but not of SOM+ interneurons, drives membrane potential towards threshold before the onset of running (Figure 5H-I), resulting in a higher firing rate during the subsequent running periods (Figure 5G). Thus, a reduction of PV+ interneuron activity during running periods in trained animals can explain several key aspects of our data. Studies in other brain regions support our conclusions: in the barrel cortex, the firing of PV+ cells dominates during quiet wakefulness (Gentet et al., 2010). Similarly, in other neocortical regions, a reduction of PV+ IN activity has been shown to affect membrane potential dynamics and neuronal firing during running states (Polack et al., 2013; Schneider et al., 2014). By contrast, inactivation of SOM+ cells leads to more heterogeneous effects on principal neurons: overall, membrane potential and firing rates during movement show less changes than during PV+ interneuron inactivation (Figure 5D-E, 5G-I and 6C-D).

### Disinhibitory role of SOM+ interneurons in shaping membrane potential ramps

PV+ interneurons densely target the perisomatic domains of principal neurons across cortical areas and layers (Packer and Yuste, 2011). By contrast, SOM+ cells densely target the tuft, apical and basal dendrites of principal neurons in a layer-specific manner (Fino and Yuste, 2011; Wang et al., 2004). Converging evidence suggests that PV+ cells strongly inhibit each other without inhibiting other interneuron subtypes, whereas SOM+ strongly inhibit PV+ interneurons across all layers without inhibiting themselves (Ährlund-Richter et al., 2019; Cottam et al., 2013; Pfeffer et al., 2013). Such a pattern of connectivity might define their distinct roles in shaping neural dynamics in the MOs neuronal network: a general and unimodal inhibitory role for PV+ interneurons, and a specific and cross-modal role in experience-dependent plasticity for SOM+ cells. For example, recent evidence has shown that such SOM+ interneuron-mediated inhibition of PV+ cells may be important to disinhibit MOs principal neurons during encoding of cue associations in an associative fear learning task (Cummings and Clem, 2020), and to synchronize network activity in the prefrontal cortex during fear expression and social discrimination (Courtin et al., 2014; Scheggia et al., 2020). If activation of SOM+ cells was exclusively and unimodally providing dendritic inhibition to principal neurons, their inhibition should result in depolarization of excitatory neurons. By contrast, we observe that inactivation of SOM+ interneurons abolishes the preparatory membrane potential ramp without affecting the mean baseline membrane potential (Figure 6B and 6D). We therefore suggest that SOM+ cells disinhibit principal neurons via inhibition of PV+ interneurons, resulting in a slow depolarizing membrane potential ramp during motor planning (Figure 7). In agreement with our suggestion, other studies have shown that during maintenance of working memory, the activity of PV+ interneurons in medial prefrontal cortex is reduced during delay periods and strongly inhibited during reward-taking periods. By contrast, SOM+ cells show a strong activation during delay periods (Kim et al., 2016a). The role of additional types of interneurons, such as vasoactive intestinal peptide (VIP) expressing cells, shown to act mainly through indirect disinhibition of principal neurons via inhibition of SOM+ and PV+ cells (Koukouli et al., 2017; Lee et al., 2013; Pi et al., 2013), remains to be explored.

### Temporal dynamics of membrane potential ramp signals during preparation of movement

As neural transmission within isolated neurons and circuits shows intrinsic time constants on the scale of milliseconds, how can preparatory activity persist during several seconds? Experiments and computational modelling have shown that sustained preparatory activity can result from neuronal modules that integrate transient inputs (Murakami et al., 2014). The robustness of the preparatory activity to large transient perturbations furthermore suggests that the network dynamics of these integrator modules in the cerebellar-thalamic-frontal network is independent, redundant and coupled through feedback connections (Chaudhuri and Fiete, 2016; Economo et al., 2018; Gao et al., 2018; Inagaki et al., 2019; Li et al., 2016; Murakami et al., 2014). Therefore, preparatory signals are thought to be initiated in frontal motor regions, from where they enter the loop and evolve during several seconds preceding motor actions (Churchland et al., 2010b; Gao et al., 2018; Li et al., 2015), regardless of how movements are initiated (Lara et al., 2018). Together, these studies propose that multi-circuit mechanisms for maintaining preparatory activity underlie the role of motor-associated cortices in the temporal organization of motor behaviors (Svoboda and Li, 2018). Consistently, our computational modelling shows that recurrent connectivity of MOs principal neurons, in concert with inhibition from PV+ and SOM+ interneurons, can convert a step-like external input into a slowly increasing and sustained ramp of activity (Figure 4 and 7).

### SOM+ plasticity can explain ramp acceleration after learning a goal-directed task

Our experiments and computational modelling suggest that PV+ and SOM+ interneurons also play a key role in producing faster membrane potential ramps with larger amplitudes during preparation of goal-driven movement (Figure 4D). Inactivation of PV+ interneurons during spontaneous movement results in depolarized membrane potential ramps with larger amplitudes, thereby reproducing some of the features of the ramps after goal-directed training. However, PV+ inactivation fails to reproduce the ramp acceleration that we observe during preparation of goal-directed movements. To fully capture all dynamics of preparatory signals after goal-directed training, we therefore suggest that excitatory inputs to SOM+ interneurons undergo task-specific experience-dependent plasticity (Biane et al., 2016). In agreement with this suggestion, other studies have shown that excitatory synaptic inputs targeting prefrontal SOM+ cells are potentiated after cue fear acquisition, thereby boosting their efficacy of disinhibition of principal neurons via potent inhibition of PV+ cells (Cummings and Clem, 2020). A similar mechanism could explain our observations: our modelling suggests that after goal-directed tasks, potentiated activity of SOM+ might exert more efficient inhibition of PV+ interneurons during preparation of movement, resulting in faster and larger depolarizing membrane potential ramps (Figure 7D).

Recent evidence suggests that MOs acts as a multisensory integration hub for adaptive choice behavior (Barthas and Kwan, 2017). The depolarising ramp that we observe may represent integration of multisensory information predicted by a recent model of the MOs circuit (Coen et al., 2021). Our whole-cell recordings, which sample from neurons without any bias for their firing rates, reveal that the synaptic integrative processes underlying the depolarizing ramp may occur at a much slower rate than was previously expected, in particular preceding spontaneous running periods. Such slow integration processes may have important implications for the synaptic and circuit basis of decision making.

## METHODS

### Animals

All procedures were carried out in accordance with the national guidelines on the ethical use of animals of EU Directive 2010/63/EU and were approved by the Ethics Committee CETEA of the Institut Pasteur (protocol number 160066). 6-to 12-week-old wild-type (WT) C57BL/6J and transgenic mice were maintained at our animal facility on a regular 12/12h light-dark cycle with ad libitum access to food and water. The following Cre mouse lines were obtained from the Jackson Laboratories: SST-Cre (no: 013044) and PV-Cre (no: 008069). These mice were backcrossed onto a C57BL/6J background. Stereotaxic injections were performed in 6-to 8-week-old male mice.

### Surgical procedures and viral vector transduction

Surgeries were performed under continuous anesthesia with isoflurane (5% for induction, 1–3% for maintenance, vol/vol). Preceding the surgery, mice were treated with buprenorphine (0.1 mg/kg i.p.) and lidocaine (0.4 mL/kg of a 1% solution, local application). Mice were positioned in a stereotaxic apparatus (David Kopf Instruments, Tujunga, CA). A half-circle stainless steel headpost (Luigs & Neumann) was fixed to the mouse skull using dental cement (Super-Bond, Sun Medical Co. Lt). Animals were allowed to recover for 2 weeks after head post implantation. Body temperature was monitored and maintained at 37°C by placing the animals on a heating pad during and after the surgery. Animals were treated with metacam (1mg/kg i.p.) before returning them to their home cages.

Circular craniotomies (0.5 mm diameter) were performed above the MOs under isoflurane anesthesia 1h before the onset of recordings using a dental drill (stereotaxic coordinates from Bregma, anteroposterior [AP] +2.7-3.1 mm, mediolateral [ML] ± 0.4-1.0 mm). Animals were treated with an injection of metacam (1mg/kg i.p.) at the end of the procedure, and then transferred to the recording setup.

To suppress the activity of PV+ or SOM+ interneurons, an adeno-associated viral vector (AAV5-hSyn-DIO-hM4D(Gi)-mCherry, ref Addgene-44362, 7E12 vector genomes (vg)/ml) was injected into the MOs of either PV-Cre or SOM-Cre mice. Another adeno-associated viral vector (AAV1-CAG-FLEX-tdTomato-WPRE, ref Addgene-28306, 1E13 vector genomes (vg)/ml) and (AAV1-CAG-tdTomato-WPRE, ref Addgene-59462, 5E12 vector genomes (vg)/ml) were used as control viruses. 6-to 8-week-old mice were injected with vectors (300~400 nL per site) into the MOs region (stereotaxic coordinates from Bregma, anteroposterior [AP] +2.7~3.1 mm, mediolateral [ML] ± 0.4~0.6 mm, 2 injections at 300 µm and 500 µm depth from dura). The virus was bilaterally pressure-injected through glass pipettes (Drummond Wiretrol 10µl) using an oil-hydraulic micromanipulator (MO-10, Narishige, Japan) at a rate of 100 nL/min. Headpost implantation was performed 3 weeks after the injections. Before the recordings were performed, CNO (Tocris Biosciences; 5 mg/kg i.p.) was administered to activate the hM4D receptor 30 min prior to recordings (Jackson et al., 2018). All recordings were performed within 3 hours after CNO injection.

### Behavioral training and analysis

Two weeks after the headpost implantation, mice were handled 10 minutes per day for 3 days. After this period, head-fixed mice were placed on a cylindrical polystyrene treadmill (20 cm diameter) supported by pressurized air bearings. Cylinder rotation associated with animal locomotion was read out from the surface of the treadmill with a computer mouse (G700s, Logitech, used in wired mode) at a poll rate of 1 kHz. As shown in Figure 1A, all mice were habituated to the treadmill 10-20 min per day for 2 consecutive days. Following habituation, mice were trained 30-45 min per day for 2-3 days to perform self-paced voluntary movement on the treadmill in a dark environment.

Another group of mice was trained in a goal-directed task in a virtual-reality environment. The virtual reality setup was implemented as described previously (Schmidt-Hieber and Häusser, 2013). Briefly, motion on the treadmill was read out as described above and linearly converted to one-dimensional movement along the virtual reality corridor. The virtual environment was projected onto a spherical dome screen (120 cm diameter), covering nearly the entire field of view of the animal, using a quarter-sphere mirror (45 cm diameter) and a projector (Casio XJ-A256) located below the mouse. The virtual linear corridor was 1.2 m long, enriched with objects placed along the linear track and vertical or oblique grating textures on the walls. A reward zone was located at the end (1.0-1.1 m) of the corridor. The Blender Game Engine (http://www.blender.org) was used in conjunction with the Blender Python API to drive the virtual reality system. Controlled water delivery was used to improve animal motivation during the goal-directed task. At the beginning of experiments, mice were placed under controlled water supply (0.5 mg of hydrogel per day, Clear H2O, BioService) and maintained at ~85% of their initial body weight over the course of behavioral training and electrophysiology experiments. The welfare and weight of animals were checked and documented on a daily basis. After habituation and water deprivation, mice underwent 6 training sessions, 30-45 min each, over the course of 1 week before recordings (Figure 1A-C). A drop of sugar water (10 μl, 8 mg/mL sucrose) was dispensed by a spout as a reward if they spent 2 s or more within the reward zone. When the animals reached the end of the linear track, they were “teleported” back to the start of the virtual corridor after crossing a black frontal wall, indicating the end of a lap and the onset of the subsequent one. Behavioral performance of the training group was comparable between different sessions (Figure 1A-C). Locomotor behavior was comparable between the trained group and the spontaneously running group (Figure 3B-D).

To detect whisker and body motion (Figure S4), we filmed the animal using an infrared-sensitive camera (Point Grey CM3-U3-13Y3M-CS) operating at a frame rate of 200 Hz. The animal was illuminated by an array of infrared LEDs. Whisker movements were analyzed with the markerless pose estimation software DeepLabCut (Mathis et al., 2018). The bases and tips of 4 whiskers, i.e. a total of 8 labels, were identified across all captured frames. To quantify whisker movement, across all captured frames we computed the sum of the Euclidean distances, in units of pixels, that were covered by each of the 8 whisker labels between adjacent frames. A whisker motion index was obtained by z-scoring the data, subtracting any offset, and low-pass filtering at *f*_c_ ~0.1 Hz. Onset of whisker movement preceding onset of running were detected where the whisker motion index continuously exceeded 10% of its maximal value within a time window of 15 s before running onset.

### Cannula implantation for muscimol Infusion

To infuse muscimol into MOs, two stainless steel guide cannulae (26 gauge; PlasticsOne, Roanoke, VA) were bilaterally implanted above the MOs (from Bregma position, anteroposterior [AP] +2.6–3.1mm, mediolateral [ML] ±0.8–1.3mm, angled at 25-30° towards medial from vertical; dorsoventral, 0.7–0.8 mm). Cannulae were anchored to the skull with dental cement (Super-Bond, Sun Medical Co. Lt). Body temperature was monitored and maintained at 37°C by placing the animals on a heating pad during and after the surgery, and the guides were covered with a dummy cannula to reduce the risk of infection. Mice were allowed to recover during 3-4 weeks from surgery before the start of water restriction, and their well-being and weight were assessed on a daily basis.

An infusion cannula (33 gauge; connected to a 1 mL Hamilton syringe via polyethylene tubing) was inserted through the guide cannula, protruding 0.5 mm, to target the MOs. 2×350 nL of Muscimol (0.6µg/µL in saline) were infused bilaterally at a rate of 100nL per min using a motorized pump (Legato 100, Kd Scientific Inc., Hilliston, MA), 45–60min before behavioral testing. To allow for penetration of the drug, the injector was maintained in position for an additional 3 min after the end of the infusion. Mice were placed back in their home cages at the end of the injection procedure.

To analyze the location and extent of the injections, we injected the fluorophore BODIPY TMR-X into MOs (Invitrogen; 5 mM in PBS 0.1 M, DMSO 40%). After 3 hours, animals were deeply anaesthetized and brains were fixed by intracardiac perfusion with paraformaldehyde (4% in PBS). Slices (50 µm) were cut with a vibratome and imaged using a confocal microscope (Opterra, Bruker). Mice were considered for further analysis if fluorescence signals could be confirmed in MOs (Figure 1F).

### *In vivo* patch-clamp electrophysiology

Whole-cell patch-clamp recordings were performed from head-fixed mice placed on the treadmill as described above. Glass pipettes were pulled from borosilicate glass (~5 MΩ pipette resistance) and filled with internal solution containing (in mM) 130 potassium methanesulphonate, 7.0 KCl, 0.3 MgCl_2_, 0.1 EGTA, 10 HEPES, 1 sodium phosphocreatine, 3.0 Na_2_ATP, 0.3 NaGTP. 5 mg/ml biocytin was added to the internal solution for staining purposes. pH was adjusted to 7.2 with KOH. Osmolarity was 289 mOsm. Whole-cell patch-clamp recordings were obtained using a standard blind-patch approach (Margrie et al., 2002; Schmidt-Hieber and Häusser, 2013). In brief, a high positive air pressure (~1000 psi) was applied to the pipettes before slowly lowering them into the dorsal part of the MOs region (Fig. 1B) via a small craniotomy (~500 µm) using a micromanipulator (Luigs & Neumann Mini In Vivo). Recordings were obtained at a depth of 150-420 µm (superficial neurons; typically layers 2/3) or 430-850 µm (deep neurons; typically layers 4–6) from the pial surface (Figure 2D). At a depth of ~150 µm from the brain surface, the air pressure was decreased to 50~80 psi. Seal resistances were always >>1 GΩ, and access resistances were typically 25~70 MΩ, with recordings terminated when access resistance exceeded 100 MΩ. Recordings were made in current-clamp mode, and no holding current was applied during recordings.

Membrane potential was low-pass filtered at 10 kHz and acquired at 50 kHz (Intan Technologies CLAMP system). During recordings, a silver/silver chloride reference electrode (0.2~0.3 mm diameter) was positioned in an additional small craniotomy close to lambda. An external solution containing (in mM) 150 NaCl, 2.5 KCl, 10 HEPES, 2 CaCl_2_, and 1 MgCl_2_ (pH 7.2, 289 mOsm) was perfused on top of the craniotomy through a round plastic chamber (4mm diameter).

### *In vivo* extracellular electrophysiology

Extracellular recordings of population activity were made using Neuropixels probes (Jun et al., 2017). Probes were lowered into the dorsal part of the MOs region to a depth of ~2.0mm measured from the brain surface via a small craniotomy (~500 µm) at an angle of 15° from vertical using a micromanipulator (Sensapex uMp-4). A silver/silver chloride wire, which was soldered to the external reference of the probes and connected to ground, was positioned in an additional small craniotomy. Recordings were performed from head-fixed mice placed on the treadmill as described above. Electrophysiological data were recorded with SpikeGLX (https://billkarsh.github.io/SpikeGLX/) and signals from the AP channels (0.3-10 kHz bandwidth, sampling rate of 30 kHz) were used for further processing. Spike sorting for unit identification was performed with Kilosort 2 (https://github.com/MouseLand/Kilosort). Identified units were manually curated using Phy (https://github.com/cortex-lab/phy). Putative cell types were assessed based on peak-to-valley ratio and half-valley width of the spike waveforms for each neuron by fitting a Gaussian mixture model (Kim et al., 2016b; Stark et al., 2013). Units with low classification confidence (P < 0.95) were unassigned. The interneuron population was further split based on average firing rate: interneurons with a mean firing rate > 10 Hz were classified as fast-spiking interneurons.

### Immunohistochemistry and cell identification

At the end of some recordings, mice were deeply anesthetized with an overdose of ketamine/xylazine (100mg/kg and 10mg/kg i.p.) and quickly perfused transcardially with 0.1 M phosphate-buffered saline followed by a 40 mg/ml PFA solution. Brains were removed from the skull and kept in PFA for at least 24 h. We stained 50-μm-thick parasagittal slices with Alexa Fluor 488–streptavidin to reveal biocytin-filled neurons and patch pipette tracts. We identified neurons as principal cells according to their characteristic electrophysiological signature (Figure 2G), including the presence of frequency adaptation during spike trains, and absence of pronounced after hyperpolarizations following action potentials (Zhao et al., 2016). Whenever the morphological recovery of recorded neurons was successful, we confirmed this classification using the shape and position of biocytin-filled neurons. In addition, the pipette tract was confirmed to terminate in MOs (Figure 2C).

### *In vivo* whole-cell electrophysiology data analysis

Input resistance was calculated from the steady-state voltage response to a small hyperpolarizing 500-ms current pulse from baseline membrane potential (Figure 2G). Only data from mice that were resting during this period were used (speed < 0.5 cm/s). Baseline membrane potential was measured before current pulse injections at the beginning of the recording. Spontaneous firing rate and membrane potential were measured across recordings with durations exceeding 60s. To analyze subthreshold membrane potential and its variance, traces were digitally low-pass filtered at 5 kHz and resampled at 10 kHz. Action potentials were then removed by thresholding to determine action potential times and then masking values 2 ms before and 10-20 ms after the action potential peak. Membrane potential oscillations were analyzed by bandpass-filtering membrane potential traces after removal of action potentials. Power spectra were computed from subthreshold membrane potential traces using Hanning windowing over data windows of 215 sampling points (~328 ms). Membrane potential variance time series were computed in rolling time windows with a width of 1000 ms.

Changes in subthreshold membrane potential (Δ*V*_m_) were computed by subtracting the mean of subthreshold membrane potential traces after spike removal (see above). To compute movement-aligned mean traces of Δ*V*_m_ (Figures 4, 6 and S6-7), we aligned traces to movement periods spanning 15 seconds before movement onset until 5 seconds after movement onset. After alignment, we computed the mean of each trace between 15 and 10 seconds before the onset of movement, and subtracted this baseline from each trace.

### Data inclusion criteria

For the analysis of intrinsic membrane properties (Figure 2E-G), cells that met basic recording criteria (initial access resistance < 70 MΩ, initial baseline membrane potential < –50 mV) from the control group of mice were included (47 neurons). For the analysis of firing rate and *V*_m_ dynamics (Figure 3-6 and S5-7), only cells with recording durations exceeding 60 s were included. Typical recordings with animals resting and running on the treadmill lasted 5–10 min, and longer recordings (~30 min) were occasionally achieved.

Running periods were defined as time intervals when the running speed continuously exceeded 1 cm/s during at least 2s. This criterion was confirmed by visual inspection of each recording to avoid inclusion of short or fractionated running periods. When we aligned data to the onset of movement (Figures 4, 6 and S6-7), running periods had to be preceded by resting periods with a duration >10s to ensure that the preparatory period was not contaminated by movement. Furthermore, the recording duration after movement onset had to exceed 5s. These criteria were applied to 29 recordings during spontaneous movement and 18 recordings during goal-directed movement, reducing the number of recordings to 23 (12 superficial and 11 deep recordings) for the control group, and 12 (5 superficial and 7 deep recordings) for the trained group of animals (Figure S6). In our analysis of preparatory spike firing (Figure 4 and 6), additional criteria were applied as we only selected neurons that spontaneously fired spikes during the movement-aligned time period. This selection further reduced the number of recordings to 11 out of 23 during spontaneous movement, and 10 out of 12 during goal-directed movement. In Figure 6, we selected recordings with resting periods >6 s preceding movement onset, and recording durations >5 s after movement onset. This selection further reduced the number of recordings to 8 out of 11 during PV+ interneurons inactivation, and 7 out of 11 during SOM+ interneurons inactivation.

Spike rates in whole-cell recordings were computed from the time points of action potential peaks (see above) in time bins of 500ms, and filtered using a sliding average with a width of 3s. Shuffled spike rate data were obtained by generating 100 artificial data sets. Each artificial data set was created from the original data set by shifting it circularly by a random number of sampling points. To quantify membrane potential ramps, we computed the mean of Δ*V*_m_ values for each recording in time bins of 1 s in movement-aligned Δ*V*_m_ traces (see above). Across all binned values of all recordings, we then computed the Spearman rank correlation to test for significant monotonous increases in membrane potential. To quantify the temporal dynamics of membrane potential and spike ramps, Spearman rank correlations were computed for different time periods during preparation of movement and during movement, as indicated in the figures and figure legends (Figure 4, 6 and S6-7).

### Quantification and statistical analysis

Wilcoxon signed-rank or Mann-Whitney U tests were used to assess the statistical significance of paired or unpaired data as appropriate. For multiple comparisons, we performed Kruskal-Wallis tests and adjusted using Dunn’s correction. In Figure 2G, S6F, S6I, S7C and S7D, a two-way ANOVA with factors’ interactions and Bonferroni post-hoc tests were used. Spearman rank correlations were computed to assess the significance of ramping dynamics (see above; Figure 4, 6 and S6-7). Statistics details are listed in Table 1. Tests were considered significant if the p value was <0.05, otherwise “n.s.” denotes “not significant”. Bar graphs and error bars show means ± s.e.m..

### Computational modelling

To simulate neuronal activity during preparation of movement, we developed a reduced model of the local MOs circuit consisting of 3 neurons: a SOM+ cell, a PV+ cell, and a principal neuron. The dynamics of the rate-based model neurons was defined by

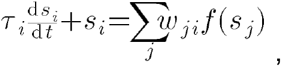

where τ _*i*_ is the time constant of neuronal integration of neuron *i, s*_*i*_ and *s*_*j*_ are the activities of neurons *i* and *j*, d*t* is the simulation time step, and *w*_*ji*_ is the synaptic weight of inputs from neuron *j* to neuron *i. f*(*x*) is a threshold function: *f*(*x*) = *x* for *x* > 0, and *f*(*x*) = 0 otherwise. All 3 neurons received input from an external source representing thalamocortical inputs. Weights of synaptic connections between neurons were adjusted to reflect our experimental results (see Supplementary Table 2). Time constants for neuronal integration τ _*i*_were set to 20 ms for both interneurons, and to 40 ms for the principal neuron. The simulation time step was set to 0.5 ms. Preparatory activity preceding the onset of running (*t* = 0s) was simulated by increasing the activity of the external inputs in a step-like fashion at *t* = –10s. Chemogenetic inactivation of PV+ or SOM+ interneurons were simulated by fixing the activities of the corresponding model neurons to 0 throughout the simulation. Plasticity of SOM+ interneurons was simulated by increasing the excitatory synaptic weights to the SOM+ model interneuron (see Supplementary Table 2).

## Acknowledgments

We thank Ian Duguid and Josh Dacre for comments on the manuscript. We thank Lucile Le Chevalier-Sontag and Claire Lecestre for their technical support. This work was supported by grants from the ERC (StG 678790 NEWRON to C.S.-H., MSCA 800027 FindMEMO to M.A., and Human Brain Project SGA2 785907 to J.-P.C. and F.K.), the Pasteur Weizmann Council, and a Pasteur-Roux fellowship to M.A.

## Notes

### Competing Interest Statement

The authors have declared no competing interest.

