## Supplemental Material for "Inhibitory control of synaptic signals preceding motor action in mouse frontal cortex"

Figure S1

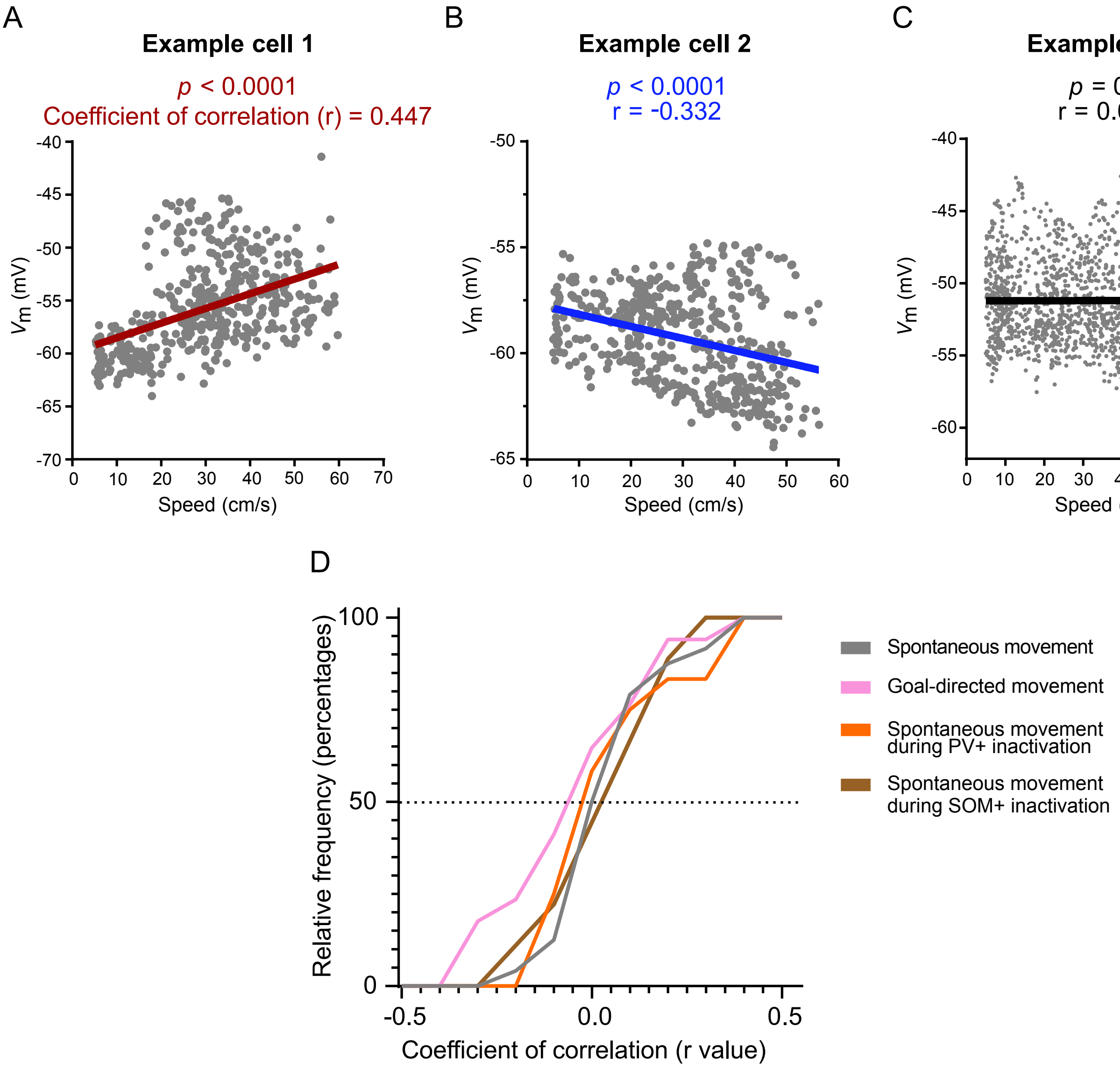


Figure S1 | Heterogeneous relationship between membrane potential and animal speed.

(A) Example recording with positive correlation between membrane potential and speed.
(B) Example recording with negative correlation between membrane potential and speed.
(C) Example recording without significant correlation between membrane potential and speed.
(D) Distribution of membrane potential – speed correlations, indicating that different types of relationships were similarly distributed across recordings.

Figure S2

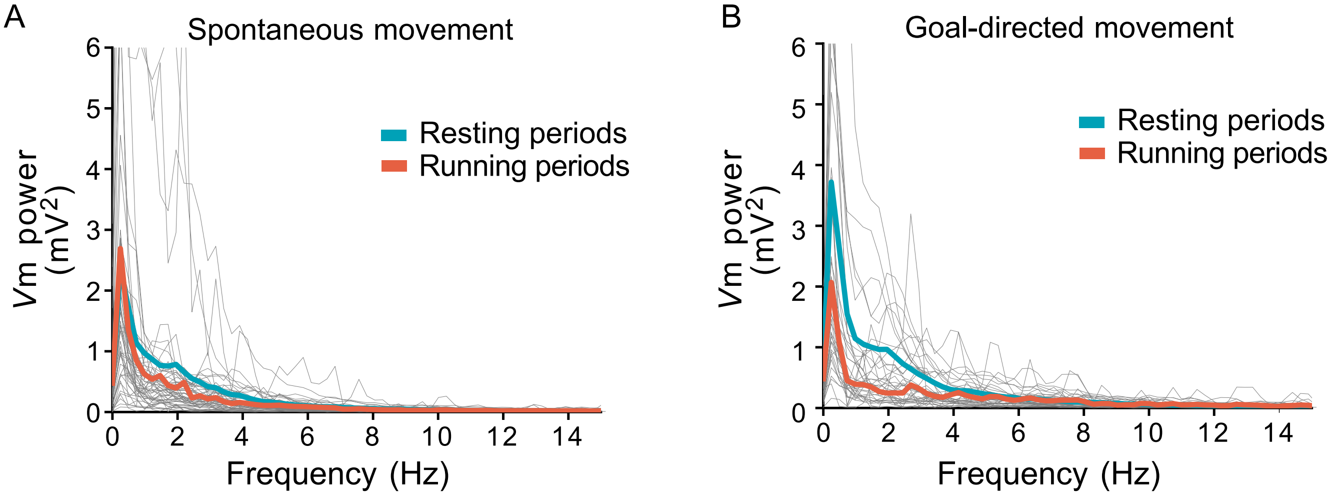


Figure S2 | Membrane potential fluctuations contain broadband frequency components without any obvious peak in the spectrum.
(A) Power spectrum density for recordings from animals running spontaneously. Grey lines represent individual recordings. Thick lines represent the means across recordings during resting periods (blue) and running periods (red).
(B) Same as in (A) for recordings from animals trained in a goal-directed task.

Figure S3

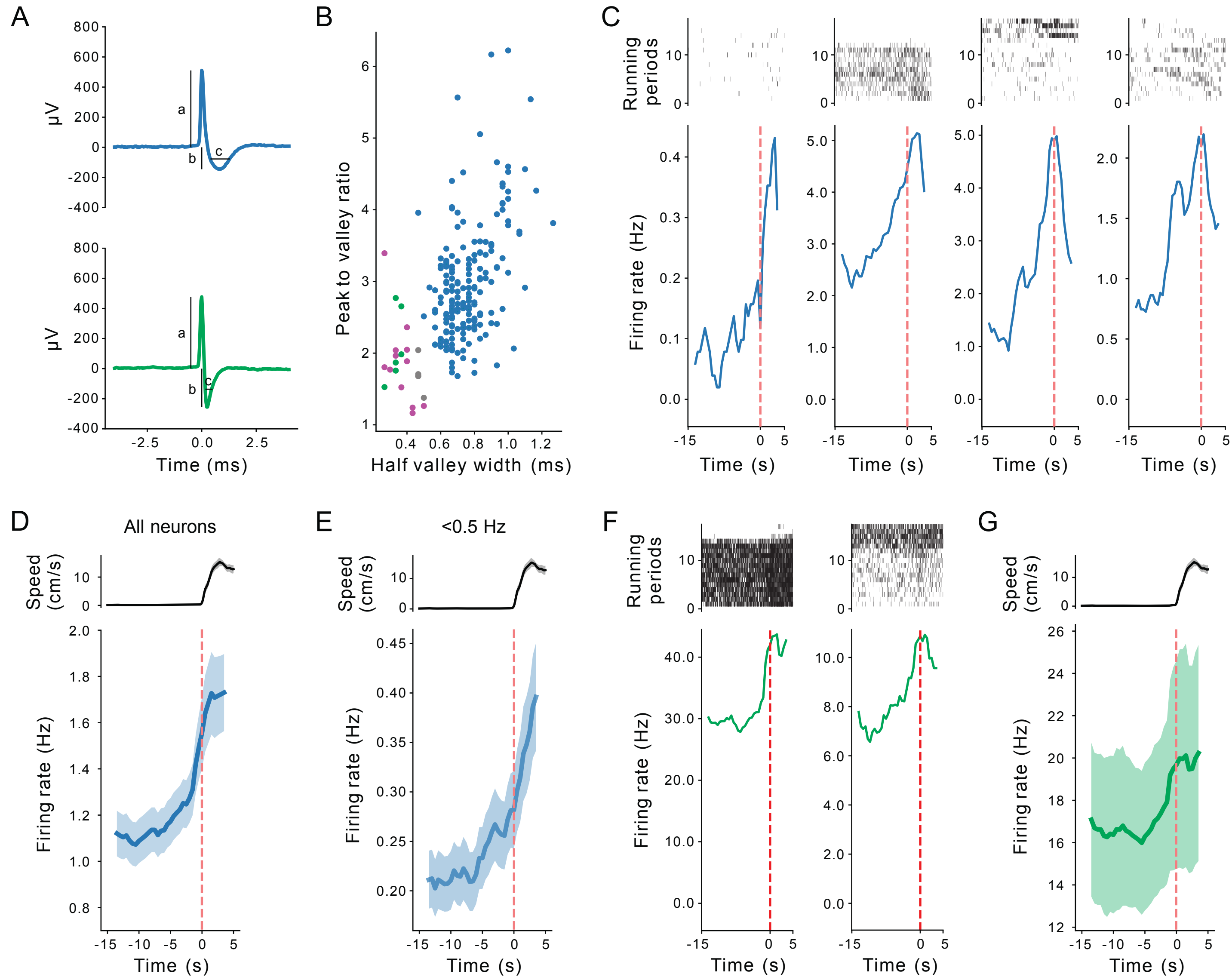


Figure S3 | Extracellular recordings of population activity from MOs of spontaneously running animals.

(A) Example spike waveforms obtained from a putative principal neuron (top) and from a putative interneuron (bottom). Traces represent mean of all action potentials produced by this unit after spike sorting. Parameters used for cell identification are indicated (a: peak; b: valley; c: half valley width).

(B) Cell type identification was performed by fitting a gaussian mixture model to the units based on the peak-to-valley ratio against half valley width. Circles correspond to individual neurons and are color-coded by putative type. Blue: principal neurons; green: fast-spiking interneurons; violet: slow-spiking interneurons; grey: low confidence assignment. Violet and grey were not used for further analysis.

(C) Example recordings from individual putative principal neurons. Data were aligned to onset of running periods at t = 0s (red dashed line). Top graphs show spike rasters for the neuron over several running periods. Bottom traces show mean firing rates for the neuron across all running periods (N = 17 periods).

(D) Mean firing rates across all putative principal neurons (N = 182 neurons) aligned to onset of running periods at t = 0s. Top graph shows mean speed across all running periods. Shaded regions indicate ± s.e.m.. Note slow ramp-up of spiking at ~t = –10s.

(E) Mean firing rates across low firing rates putative principal neurons (< 0.5 Hz, N = 66 neurons) aligned to onset of running periods at t = 0s.  Shaded regions indicate ± s.e.m.. Note slow ramp-up of spiking at ~t = –10s.

(F) Example recordings from individual putative fast-spiking interneurons, presented as examples in (C).

(G) Mean firing rates across all putative interneurons aligned to onset of running periods at t = 0s. Top graph shows mean speed across all running periods. Shaded regions indicate ± s.e.m.. Note slow ramp-up of spiking at ~t = –5s.

Figure S4

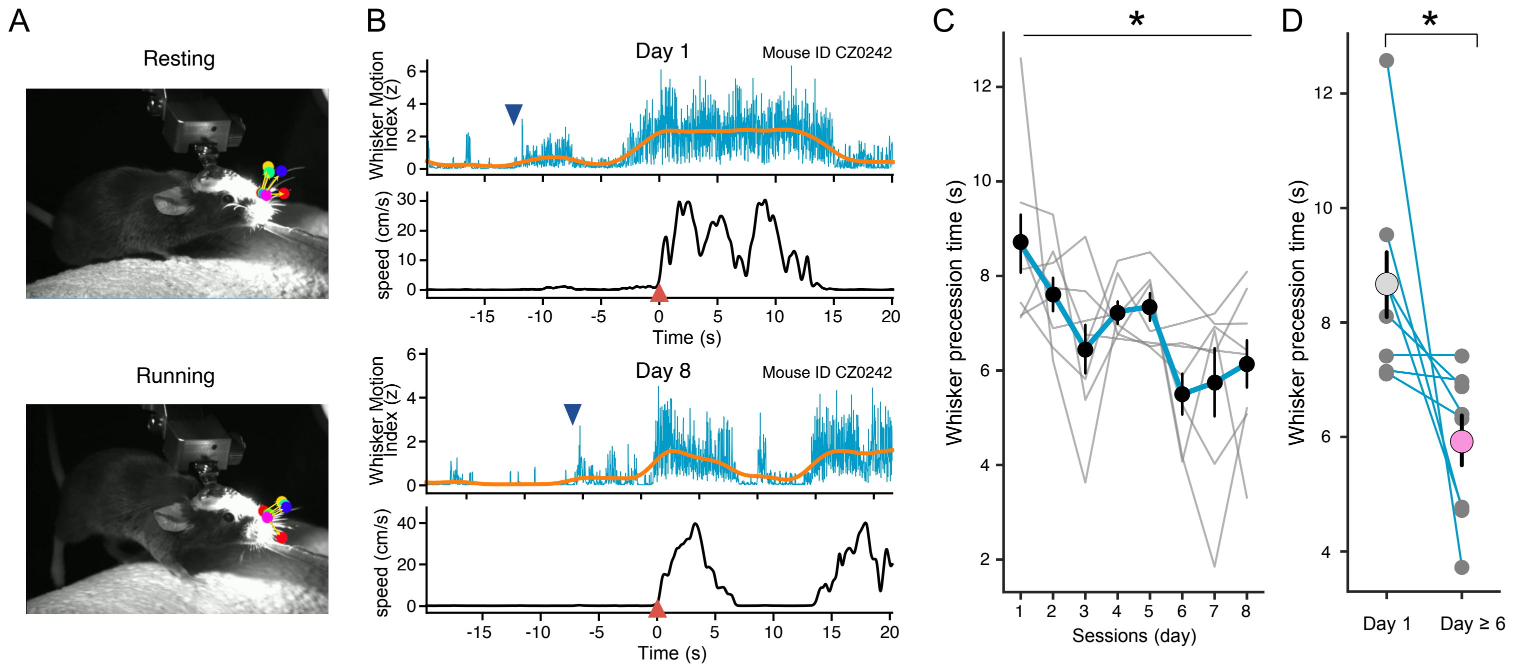


Figure S4 | Whisker motion precedes running onset.

(A) Example frames of an infrared-illuminated animal during resting (top) and running states (bottom). Filled circles indicate labels (whisker bases and tips) identified by DeepLabCut.

(B) Whisker motion index and animal speed are plotted against time during two example running periods. Blue traces represent raw data, red traces represent low-pass filtered data. The blue triangle indicates onset of whisker movement, the red triangle indicates running onset (t = 0s).

(C) Whisker precession time (i.e. the difference between the times indicated by the red and blue arrows in B) is plotted against training session number. Grey traces represent individual animals, blue trace and black symbols represent the mean across animals.

(D) Comparison of whisker precession times between the first session (8.68 ± 0.68 s) and the mean of sessions on day 6 or later (5.94 ± 0.50 s, *n* = 8). Small symbols and lines represent individual animals, large symbols represent mean across animals.

Error bars represent s.e.m. Statistical significance was assessed using repeated measures ANOVA (C), and Wilcoxon signed rank tests (D) for day 1 vs day 6. ns, not significant; *$p<0.05$. Mean values, s.e.m., and statistics details are provided in Table S1.

### Figure S5

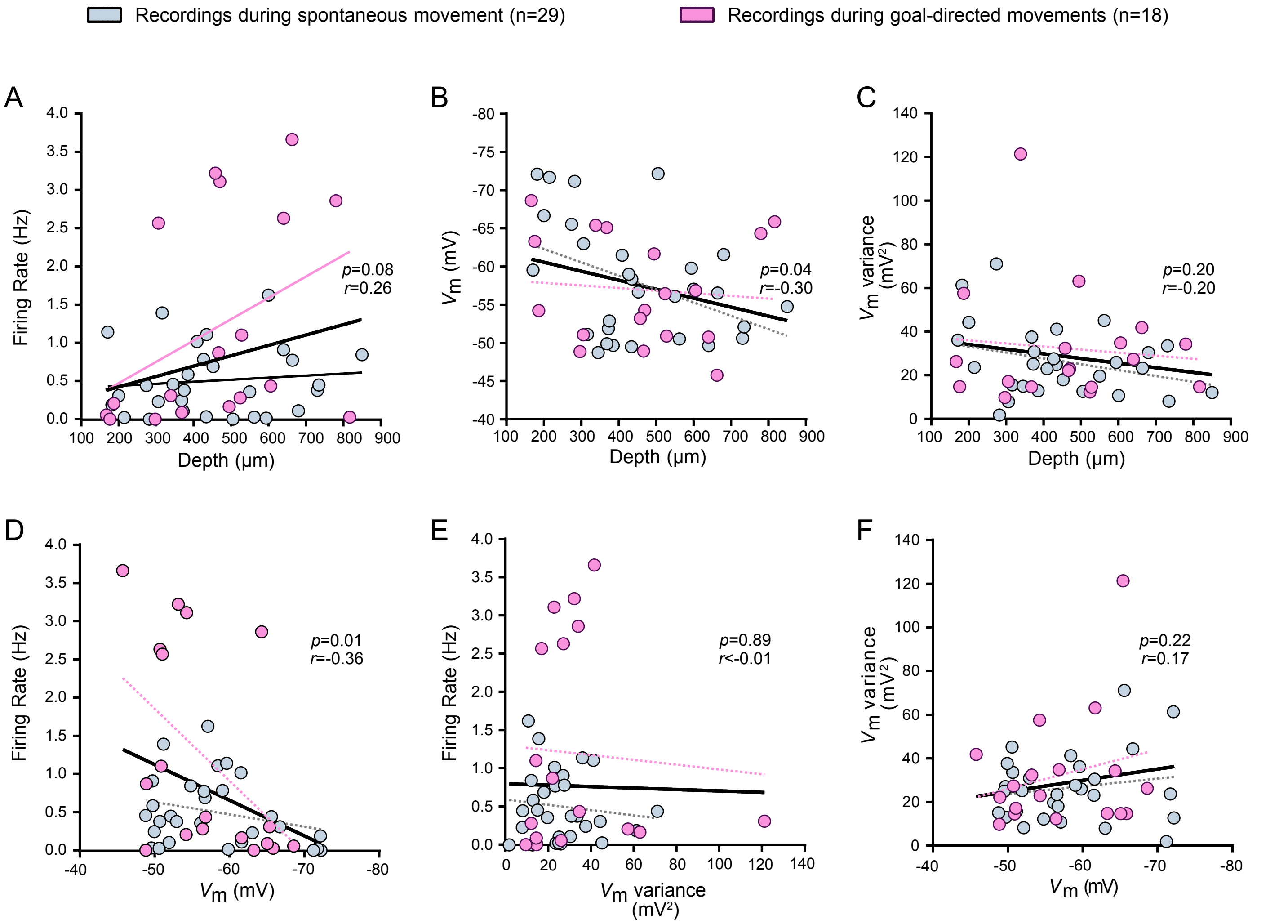


Figure S5 | Relationships between firing rates, membrane potential dynamics, and recording depth.

(A–C), Relationships between firing rate (A), baseline membrane potential (B), membrane potential variance (C) and recording depth for untrained (grey) and trained (pink) animals.
(D–F), Relationships between firing rate and membrane potential (D), firing rate and membrane potential variance (E), and membrane potential variance and membrane potential (F) for untrained (grey) and trained (pink) animals.

Statistics values and black lines refer to linear regression analysis applied to all recordings from trained and untrained animals.

### Figure S6

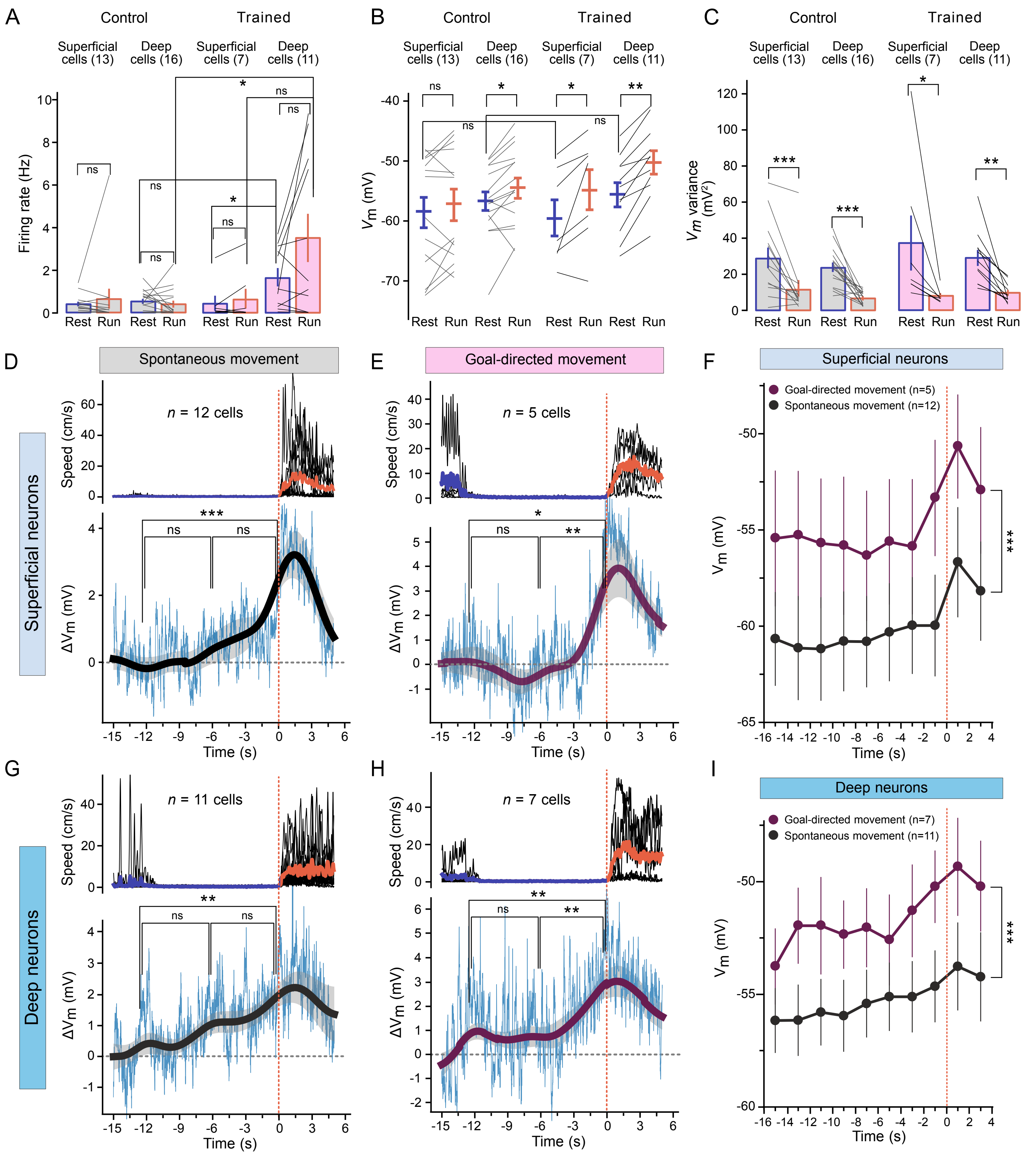
A
Control
Trained
Superficial cells (13)
Deep cells (16)
Superficial cells (7)
Deep cells (11)
Firing rate (Hz)
Rest Run Rest Run Rest Run Rest Run
B
Vm (mV)
C
Vm variance (mV2)
D
Spontaneous movement
Superficial neurons
n = 12 cells
Speed (cm/s)
ΔVm (mV)
Time (s)
E
Goal-directed movement
n = 5 cells
F
Superficial neurons
Goal-directed movement (n=5)
Spontaneous movement (n=12)
Vm (mV)
Time (s)
G
Deep neurons
n = 11 cells
H
n = 7 cells
I
Deep neurons
Goal-directed movement (n=7)
Spontaneous movement (n=11)

Figure S6 | Subthreshold membrane potential ramps precede the onset of movement in both superficial and deep neurons.

(A-C) Summary of firing rates (A; superficial neurons in control group: resting 0.42 ± 0.12 Hz *vs* running 0.67 ± 0.51 Hz, n = 13; deep neurons in control group: resting 0.57 ± 0.12 Hz *vs* running 0.44 ± 0.17 Hz, n = 16; superficial neurons in trained group: resting 0.48 ± 0.38 Hz *vs* running 0.67 ± 0.50 Hz, n = 7; deep neurons in trained group: resting 1.67 ± 0.45 Hz *vs* running 3.53 ± 1.18 Hz, n = 11) mean *V*m (B; superficial control resting -59.6 ± 2.7mV *vs* running -58.0 ± 2.7mV; deep control resting -56.7 ± 1.5mV *vs* running -54.4 ± 1.7mV; superficial trained resting -59.5 ± 3.2mV *vs* running -54.8 ± 3.7mV; deep trained resting -55.4 ± 2.0mV vs running -50.1 ± 2.0mV) and *V*m variance (C; superficial control resting 29.5 ± 5.9mV$^2$ *vs* running 12.2 ± 4.8mV$^2$; deep control resting 24.0 ± 2.7mV$^2$ *vs* running 7.2 ± 0.9mV$^2$; superficial trained resting 37.2 ± 16.5mV$^2$ *vs* running 7.9 ± 1.7mV$^2$; deep trained resting 29.0 ± 4.7mV$^2$ vs running 9.6 ± 1.7mV$^2$) during resting (blue) and running periods (red) for all recordings from the control and trained groups in superficial and deep neurons (control superficial cells at 291 ± 22 μm; control deep cells at 579 ± 34 μm; trained superficial cells at 263 ± 34 μm; trained deep cells at 586 ± 40 μm).

(D-E) Summary of membrane potential changes (Δ*V*m) aligned to the onset of running periods in superficial neurons (n=12 recordings from control animals (D), n=5 recordings from trained animals (E)). Data were aligned to the onset of running periods as shown in Fig. 4C–D. Mean *V*m between $t$ = –15 s and $t$ = –10 s was used as a baseline to calculate Δ*V*m. Top, mean animal speed. Bottom, mean membrane potential (thin traces), mean low-pass filtered membrane potential (thick traces, shaded regions represent mean ± s.e.m). Note that all spiking and non-spiking neurons which have enough long recording periods before and after onset of movements are included in the figure (see Methods).

(F) Comparison of mean *V*m in superficial neurons between the control group and the trained group during resting and running periods. Data were binned in 2-s time bins ($F$=7.846, $p$=0.0004).
(G-H) Same as in (D–E) for recordings from deep neurons (n = 11 recordings from control animals (G), and n = 7 recordings from trained animals (H)).
(**I**) Same as (F) for recordings from deep neurons ($F$=11.02, $p$<0.0001 for **I**).

For this figure, the data set shown in Figure 3 was split into recordings from deep and superficial neurons.

Error bars represent ±s.e.m. Statistical significance was assessed by Wilcoxon signed rank tests (A-C) for paired groups, and Mann-Whitney tests (A-B) for unpaired groups, Spearman's correlation (D-E and G-H) and two-way ANOVA with Bonferroni post-hoc tests (F and **I**). ns, not significant; *$p$<0.05; **$p$<0.01; ***$p$<0.001. Mean values, s.e.m., and statistics details are provided in Table S1.

Figure S7

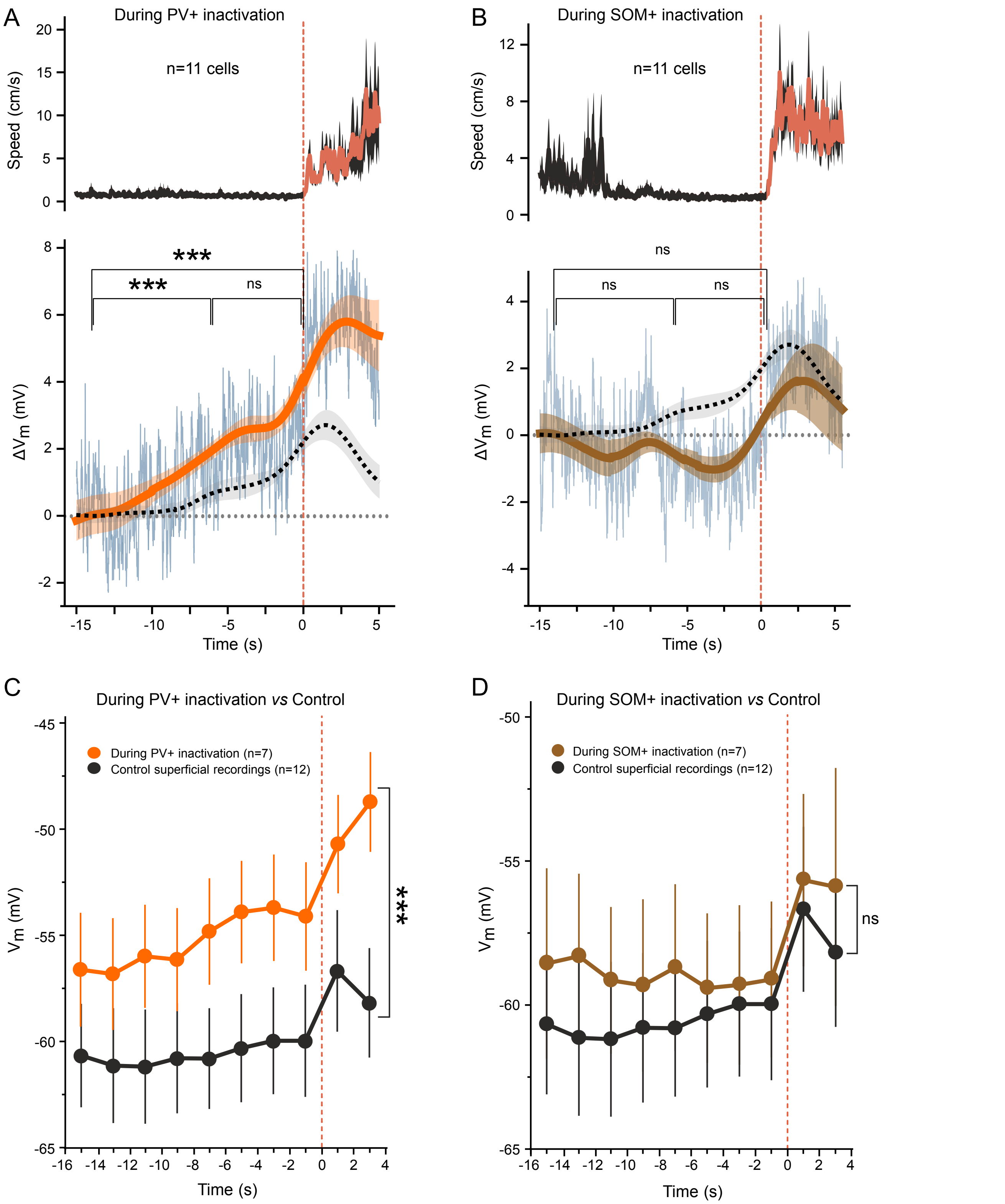


Figure S7 | Inactivation of local PV+ in MOs, but not of SOM+, affects membrane potential before and during running onset.

(A-B) Summary of speed and membrane potential of all spiking and non-spiking recordings preceding movement onset during chemogenetic inactivation of PV+ interneurons (A, n=11; 8 spiking and 3 non-spiking neurons) or of SOM+ interneurons (B, n=11; 7 spiking and 4 non-spiking neurons). Recordings from only the spiking subset of these recordings is shown in Figure 6C-D. Data are aligned to running onset at t = 0 s (indicated by a red vertical dashed line). Top, mean animal speed. Bottom, mean membrane potential (thin traces), mean low-pass filtered membrane potential (thick traces, shaded regions represent mean ± s.e.m). The black dash traces represent the control recordings including spiking and non-spiking neurons (n=23; 11 spiking and 12 non-spiking neurons).

(C-D) Comparison of mean $V_m$ in superficial neurons between control recordings (black, same as shown in Figure S6F) and recordings during PV+ inactivation (orange) aligned to the onset of running periods. Data were binned in 2-s time bins ($F$=11.10, $p$<0.0001). Same as (B) for recordings from superficial neurons during inactivation of SOM+ interneurons ($F$=0.7720, $p$=0.2454).

Error bars represent ±s.e.m. Statistical significance was assessed by Spearman's correlation (A-B) and two-way ANOVA with Bonferroni post-hoc tests (C-D). ns, not significant; *$p$<0.05; **$p$<0.01; ***$p$<0.001. Statistics details are provided in Table S1.

#### Supplemental Table 1.

| *Figure* | *Variable* | *Depth of recordings* | *Numbers of recordings* | *Values (mean±sem)* | *Statistics test* |
|---|---|---|---|---|---|
| Fig 1B | Reward rate (Reward numbers/teleportations) | None | 20 mice | Day1: 0.31±0.04; Day2: 0.35±0.05;<br>Day3: 0.38±0.05; Day4: 0.40±0.05;<br>Day5: 0.43±0.04; Day6: 0.47±0.05. | Repeated meausres ANOVA F=2.885, *p*=0.0415; Wilcoxon test for day1 vs day6, *p*=0.0006; |
| Fig 1C | Success rate<br>(Lick hits/rewards) | | 20 mice | Day1: 0.31±0.05; Day2: 0.42±0.05;<br>Day3: 0.46±0.05; Day4: 0.55±0.05;<br>Day5: 0.52±0.05; Day6: 0.55±0.05. | Repeated meausres ANOVA F=2.529, *p*=0.0340; Wilcoxon test for day1 vs day6, *p*=0.0042; |
| Fig 1G | Reward rate with Muscimol (Reward umbers/teleportations) | | 5 mice with cannula implantation | Day before: 0.46±0.14;<br>Day with Muscimol: 0.07±0.02;<br>Day after:0.37±0.17. | Repeated meausres ANOVA F=7.679, p=0.0138; |
| Fig 1H | Success rate with Muscimol<br>(Lick hits/reward) | | 5 mice with cannula implantation | Day before: 0.49±0.14;<br>Day with Muscimol: 0.07±0.07;<br>Day after:0.48±0.12. | Repeated meausres ANOVA F=4.714, p=0.0444; |
| Fig 2E | Basal membrane potential: superficial vs. deep recordings. | Superficial cells across 150~420µm:<br>Average depth 303±16µm;<br><br>Deep cells across 430~850µm:<br>Average depth 547±24µm. | Superficial cells: 24;<br><br>Deep cells: 23. | Superficial cells: -67.6±1.6mV;<br>Deep cells: -60.5±1.2mV. | Mann-Whitney test: *p*=0.0014. |
| Fig 2F | Input resistance: superficial vs. deep recordings. | | | Superficial cells: 69.9±6.8Mohm;<br>Deep cells: 77.0±7.6Mohm. | Mann-Whitney test: *p*=0.3922 |
| Fig 2G | Firing rate and current injection curve: superficial vs. deep recordings. | | | Superficial cells at 300pA injection: 14.4±2.3Hz;<br>Deep cells at 300pA injection: 21.0±2.3Hz. | two-way ANOVA with Bonferroni post-hoc tests; comparision of different depths: F=9.94, *p*=0.002; comparision of two groups at 300pA, *p*=0.01. |
| Fig 3B | Mean speed (cm/s) | Same as Figure 3G-J<br>Control group across 170~850µm;<br><br>Control superficial group:<br>Average depth 291±22µm; Deep group: 579±34µm;<br><br>Trained group across 167~817µm;<br><br>Trained superficial group:<br>Average depth 263±34µm; Deep group: 586±40µm. | <u>Control group: 29;</u><br><br>Trained group: 18.<br>Control group: 29;<br><br>Control superficial cells: 13; Deep cells: 16.<br><br><u>Trained group: 18.</u><br><br>Trained superficial cells: 7; Deep cells: 11. | Control: 13.1 ± 1.4<br>Trained: 21.8 ± 1.7 | Mann-Whitney test: *p*=0.0002. |
| Fig 3C | Running period duration (s) | | | Control: 10.8 ± 1.4<br>Trained: 7.7 ± 0.9 | Mann-Whitney test: *p*=0.0943. |
| Fig 3D | Running frequency per minute | | | Control: 0.6 ± 0.1<br>Trained: 1.1 ± 0.3 | Mann-Whitney test: *p*=0.0204. |
| Fig 3G | Firing rate during resting vs. running. | | | Control resting: 0.50±0.08Hz;<br>Control running: 0.54±0.24Hz;<br>Trained resting: 1.21±0.33Hz;<br>Trained running: 2.42±0.80Hz. | Control resting vs. running: Wilcoxon test, *p*=0.1004; Trained resting vs. running: wilcoxon test, *p*=0.1324; Control resting vs. Trained resting: Mann-Whitney test: *p*=0.4638; Control running vs. Trained running: Mann-Whitney test: *p*=0.0760. |
| Fig 3H | Mean membrane potential during resting vs. running. | | | Control resting: -58.0±1.4mV;<br>Control running: -56.0±1.5mV;<br>Trained resting: -57.0±1.7mV;<br>Trained running: -51.9±1.8mV. | Control resting vs. running: Wilcoxon test, *p*=0.0052; Trained resting vs. running: wilcoxon test, *p*<0.0001; Control resting vs. Trained resting: Mann-Whitney test: *p*=0.7699; Control running vs. Trained running: Mann-Whitney test: *p*=0.0859. |
| Fig 3I | Delta Mean membrane potential before and after running: control vs. trained groups. | | | Control: 1.98±0.65mV;<br>Trained: 5.04±0.87mV. | Control vs. Trained: Mann-Whitney test: *p*=0.0060. |
| Fig 3J | Membrane potential Variance: resting vs. running. | | | Control resting: 26.5±3.0mV$^2$;<br>Control running: 9.5±2.2mV$^2$;<br>Trained resting: 32.5±6.5mV$^2$;<br>Trained running: 9.0±1.2 mV$^2$. | Control resting vs. running: Wilcoxon test, *p*<0.0001; Trained resting vs. running: wilcoxon test, *p*<0.0001; Control resting vs. Trained resting: Mann-Whitney test: *p*=0.7699; Control running vs. Trained running: Mann-Whitney test: *p*=0.6258. |
| Fig 4C | Membrane potential ramps at different time window. | Control group across 170~850µm;<br><br>Control superficial group:<br>Average depth 248±33µm;<br>deep group: 629±85µm; | <u>Control group: 11 spiking recordings out of 29 neurons in Figure 3G-J;</u><br><br>Control superficial cells: 6; deep cells: 5. | None | Spearman's correlation:<br>-14~-0s window, r=0.39, *p*<0.0001;<br>-14~-6s window, r=0.30, *p*=0.0048;<br>-6~0s window, r=0.20, *p*=0.1073. |
| | Firing rate ramps at different time window. | | | | Spearman's correlation:<br>-15~-0s window, r=0.2124, *p*=0.00008;<br>-6~-0s window, r=0.1188, *p*=0.1286. |
| Fig 4D | Membrane potential ramps at different time window. | Trained group across 167~817µm.<br><br>Trained superficial group:<br>Average depth 277±33µm;<br>Deep group: 548±36µm. | <u>Trained group: 10 spiking recordings out of 18 neurons in Figure 3G-J;</u><br><br>Trained superficial cells: 3; deep cells: 7. | None | Spearman's correlation:<br>-14~-0s window, r=0.31, *p*=0.0002;<br>-14~-6s window, r=0.04, *p*=0.7173;<br>-6~0s window, r=0.49, *p*=0.0001. |
| | Firing rate ramps at different time window. | | | | Spearman's correlation:<br>-15~-0s window, r=0.0147, *p*=0.7972;<br>-6~-0s window, r=0.2455, *p*=0.0025. |
| Fig 5D | Firing rate during resting periods. | Control group across 170~850µm;<br><br>Control superficial cells: 291±22µm; Deep cells: 579±34µm; | Control group: 29;<br>Superficial recordings:16;<br>Deep recordings:13. | Control group resting: 0.50±0.08Hz;<br><br>During inactivation of PV+ INs: 0.82±0.11Hz;<br><br>During inactivation of SOM+ INs: 0.63±0.17Hz; | Kruskal-Wallis test with Dunn's multiple comparisons test, p=0.0132.<br>Control vs PV+ INs inactivation: *p*=0.0238;<br>Control vs SOM+ INs inactivation: *p*=0.3671;<br>PV+ INs inactivation vs SOM+ INs inactivation: *p*=0.0066. |

| | | | | | |
|---|---|---|---|---|---|
| Fig 5E | Mena membrane potential during resting periods. | Recordings during PV+ inactivation across 169~856µm;<br><br>Superficial cells: 289±26µm; Deep cells: 557±30µm; | PV+ INs inactivation: 28; Superficial:13; Deep: 15. | Control group resting: -58.0±1.4mV;<br><br>During inactivation of PV+ INs: -58.1±1.3mV;<br><br>During inactivation of SOM+ INs: -60.6±1.1mV. | Kruskal-Wallis test with Dunn's multiple comparisons, *p*=0.1132.<br><br>Control vs PV+ INs inactivation: *p*=0.8863;<br>Control vs SOM+ INs inactivation: *p*=0.0913;<br>PV+ INs inactivation vs SOM+ INs inactivation: *p*=0.0629. |
| Fig 5F | Membrane potential variance during resting periods. | Recording during SOM+ inactivation across 190~847µm;<br><br>Superficial cells: 318±20µm; Deep cells: 622±36µm; | SOM+ INs inactivation: 34. Superficial:17; Deep:17. | Control group resting: 26.5±3.0$mV^2$;<br><br>During inactivation of PV+ INs: 45.1±4.3$mV^2$;<br><br>During inactivation of SOM+ INs: 35.6±3.6$mV^2$; | Kruskal-Wallis test with Dunn's multiple comparisons, *p*=0.0049.<br>Control vs PV+ INs inactivation: *p*=0.0012;<br>Control vs SOM+ INs inactivation: *p*=0.0522;<br>PV+ INs inactivation vs SOM+ INs inactivation: *p*=0.1510. |
| Fig 5G | Firing rate during resting and running periods. | Control group across 170~850µm;<br><br>Control superficial cells: 291±22µm; Deep cells: 579±34µm;<br><br>Recordings during PV+ inactivation across 169~856µm;<br><br>Superficial cells: 304±31µm; Deep cells: 631±40µm; | Control group: 29; Superficial cells:16; Deep cells:13.<br><br>During PV+ INs inactivation: 14; Superficial cells: 9; Deep cells: 5. | Control resting: 0.50±0.08Hz;<br>Control running: 0.54±0.24Hz;<br>PV+ INs inactivation resting: 0.86± 0.14Hz; running: 2.77±1.02Hz;<br>SOM+ INs inactivation resting: 0.71± 0.33Hz; running: 2.07±1.10Hz. | Control resting vs. running: Wilcoxon test, *p*=0.1004; PV+ INs inactivation resting vs running: Wilcoxon test, *p*=0.0785; SOM+ INs inactivation resting vs running: Wilcoxon test, *p*=0.4283;<br><br>PV+ INs inactivation resting vs Control resting: Mann-Whitney test, *p*=0.0309; PV+ INs inactivation running vs Control running: Mann-Whitney test, *p*=0.0002; SOM+ INs inactivation running vs Control running: Mann-Whitney test, *p*=0.2953. |
| Fig 5H | Mean membrane potential during resting and running periods. | | | Control resting: -58.0±1.4mV;<br>Control running: -56.0±1.5mV;<br>PV+ INs inactivation resting: -60.3± 1.4mV; running: -53.0±2.2mV;<br>SOM+ INs inactivation resting: -61.8mV±1.7mV; running: -58.3± 2.7mV; | Control resting vs. running: Wilcoxon test, *p*=0.0052; PV+ INs inactivation resting vs running: Wilcoxon test, *p*=0.0004; SOM+ INs inactivation resting vs running: Wilcoxon test, *p*=0.1294; |
| Fig 5I | Delta Mean membrane potential during resting and running periods. | Recording during SOM+ inactivation across 190~859µm;<br><br>Superficial cells: 344±34µm; Deep cells: 684±113µm; | During SOM+ INs inactivation: 12; Superficial cells: 9; Deep cells: 3. | Control: 1.98±0.65mV;<br>PV+ INs inactivation: 7.3±1.4mV;<br>SOM+ INs inactivation: 3.5±2.0mV. | Kruskal-Wallis test with Dunn's multiple comparisons, p=0.0070.<br>Control vs PV+ INs inactivation: *p*=0.0050;<br>Control vs SOM+ INs inactivation: *p*>0.9999;<br>PV+ INs inactivation vs SOM+ INs inactivation: *p*=0.1635. |
| Fig 5J | Membrane potential variance during resting and running periods. | | | Control resting: 26.5±3.0$mV^2$;<br>Control running: 9.5±2.2$mV^2$;<br>PV+ INs inactivation resting: 39.9± 7.2$mV^2$; running: 15.9±4.0$mV^2$;<br>SOM+ INs inactivation resting: 37.2± 5.7$mV^2$; running: 23.1±5.8$mV^2$. | Control resting vs. running: Wilcoxon test, *p*<0.0001; PV+ INs inactivation resting vs running: Wilcoxon test, *p*=0.0001; SOM+ INs inactivation resting vs running: Wilcoxon test, *p*=0.0161. |
| Fig 6C | Membrane potential ramps at different time window, during PV+ inactivation. | Spiking recordings during PV+ inactivation across 169~520µm;<br><br>Average depth: 365±45µm. | Control spiking recordings used in Figure 4C;<br><br>Spiking recordings during PV+ inactivation: 8 out of 14 recordings in Figure 5G-J, which meet the criteria; | None | Spearman's correlation:<br>-14~-0s window, r=0.48, *p*=0.0000;<br>-14~-6s window, r=0.50, *p*=0.0000;<br>-6~0s window, r=-0.09, *p*=0.5383. |
| | Firing rate ramps at different time window, during PV+ inactivation.. | | | | Spearman's correlation:<br>-15~-0s window, r=0.1716, *p*=0.0328;<br>-6~-0s window, r=0.2915, *p*=0.0111; |
| Fig 6D | Membrane potential ramps at different time window, during SOM+ inactivation. | Spiking recordings during SOM+ inactivation across 190~850µm;<br><br>Average depth: 492±107µm. | Control spiking recordings used in Figure 4C;<br><br>Spiking recordings during SOM+ inactivation: 7 out of 12 recordings in Figure 5G-J, which meet the criteria; | None | Spearman's correlation:<br>-14~-0s window, r=-0.13, *p*=0.2172;<br>-14~-6s window, r=-0.02, *p*=0.8906;<br>-6~0s window, r=0.03, *p*=0.8367. |
| | Firing rate ramps at different time window, during SOM+ inactivation.. | | | | Spearman's correlation:<br>-15~-0s window, r=-0.13750, p=0.0429;<br>-6~-0s window, r=0.1227, p=0.2125; |
| Fig S4C-D | Whisker precession times (second); day 1 vs day 6. | None | 8 mice | Day 1: 8.68±0.68 s;<br>Day 6: 5.94±0.50 s. | Repeated meausres ANOVA F=3.726, *p*=0.0377; Wilcoxon test for day1 vs day6, *p*=0.0234. |
| Fig S6A | Firing rate during resting vs. running. | Control superficial group:<br>Average depth 291±22µm;<br><br>Control deep group:<br>Average depth 579±34µm;<br><br>Trained superficial group:<br>Average depth 263±34µm;<br><br>Trained deep group across:<br>Average depth | Control superficial cells: 13; Deep cells:16;<br><br>Trained superficial cells: 7; Deep cells: 11. | Control surf resting: 0.42±0.12Hz;<br>Control surf running: 0.67±0.51Hz;<br><br>Control deep resting: 0.57±0.12Hz;<br>Control deep running: 0.44±0.17Hz;<br><br>Trained surf resting: 0.48±0.38Hz;<br>Trained surf running: 0.67±0.50Hz.<br><br>Trained deep resting: 1.67±0.45Hz;<br>Trained deep running: 3.53±1.18Hz. | Control superficial cells resting vs. running: Wilcoxon test, *p*=0.1099;<br>Control deep cells resting vs. running: Wilcoxon test, *p*=0.3303;<br>Trained superficial cells resting vs. running: Wilcoxon test, *p*=0.5625;<br>Trained superficial cells resting vs. running: Wilcoxon test, *p*=0.2061;<br><br>Trained superficial cells resting vs. Trained deep cells resting: Mann-Whitney test, *p*=0.0260;<br>Trained superficial cells running vs. Trained deep cells running: Mann-Whitney test, *p*=0.0932.<br><br>Control deep cells running vs. Trained deep cells running: Mann-Whitney test, *p*=0.0404.<br>Control deep cells resting vs. Trained deep cells resting: Mann-Whitney test, |

| | | | | | |
|---|---|---|---|---|---|
| | | 586±40μm. | | | $p$=0.0635. |
| Fig S6B | Membrane potential during resting vs. running. | | | Control surf resting: -59.6±2.7mV;<br>Control surf running: -58.0±2.7mV;<br><br>Control deep resting:-56.7±1.5mV;<br>Control deep running:-54.4±1.7mV;<br><br>Trained surf resting: -59.5±3.2mV;<br>Trained surf running: -54.8±3.7mV;<br><br>Trained deep resting: -55.4±2.0mV;<br>Trained deep running: -50.1±2.0mV. | Control superficial cells resting vs. running: Wilcoxon test, $p$=0.1677;<br>Control deep cells resting vs. running: Wilcoxon test, $p$=0.0182;<br>Trained superficial cells resting vs. running: Wilcoxon test, $p$=0.0313;<br>Trained superficial cells resting vs. running: Wilcoxon test, $p$=0.0010;<br>Control superficial cells resting vs. trained superficial cells resting: Mann-Whitney test, $p$=0.9385;<br>Control deep cells resting vs. Trained deep cells resting: Mann-Whitney test, $p$=0.6800. |
| Fig S6C | Membrane potential variance during resting vs. running. | | | Control surf resting: 29.5±5.9mV$^2$;<br>Control surf running: 12.2±4.8mV$^2$;<br><br>Control deep resting: 24.0±2.7mV$^2$;<br>Control deep running: 7.2±0.9mV$^2$;<br><br>Trained surf resting: 37.2±16.5mV$^2$;<br>Trained surf running: 7.9±1.7mV$^2$;<br><br>Trained deep resting: 29.0±4.7mV$^2$;<br>Trained deep running: 9.6±1.7mV$^2$. | Control superficial cells resting vs. running: Wilcoxon test, $p$=0.0005;<br>Control deep cells resting vs. running: Wilcoxon test, $p$<0.0001;<br>Trained superficial cells resting vs. running: Wilcoxon test, $p$=0.0156;<br>Trained superficial cells resting vs. running: Wilcoxon test, $p$=0.0010. |
| Fig S6D | Membrane potential ramps at different time window. | Control superficial group across 170~385μm: Average depth 284±22μm;<br><br>Trained superficial group across 167~368μm: Average depth 299±28μm; | Control superficial group: 12 cells; 6 spiking neurons and 6 non-spiking neurons during preparation of movement.<br><br>Trained superficial group: 5 cells. 3 spiking neurons and 2 non-spiking neurons. | None | Spearman's correlation:<br>-12~-0s window, r=0.28, $p$=0.0007;<br>-12~-6s window, r=0.07, $p$=0.5391;<br>-6~0s window, r=0.18, $p$=0.1277. |
| Fig S6E | Membrane potential ramps at different time window. | | | | Spearman's correlation:<br>-12~-0s window, r=0.30, $p$=0.0220;<br>-12~-6s window, r=-0.22, $p$=0.2381;<br>-6~0s window, r=0.56, $p$=0.0014. |
| Fig S6F | Mean membrane potential control superficial vs. trained superficial recordings | | | | Two way ANOVA with Bonferroni post-doc tests: between groups, F=7.846, $p$=0.0004. |
| Fig S6G | Membrane potential ramps at different time window. | Control deep group across 430~850μm: Average depth 590±34μm;<br><br>Trained deep group across 458~817μm: Average depth 548±36μm. | Control deep group: 11 cells; 6 spiking neurons and 6 non-spiking neurons.<br><br>Trained deep group: 7 cells. 7 spiking neurons. | None | Spearman's correlation:<br>-12~-0s window, r=0.27, $p$=0.0019;<br>-12~-6s window, r=0.10, $p$=0.4077;<br>-6~0s window, r=0.22, $p$=0.0758. |
| Fig S6H | Membrane potential ramps at different time window. | | | | Spearman's correlation:<br>-12~-0s window, r=0.30, $p$=0.0049;<br>-12~-6s window, r=0.03, $p$=0.8651;<br>-6~0s window, r=0.20, $p$=0.0089. |
| Fig S6I | Mean membrane potential control deep vs. trained deep recordings | | | | Two way ANOVA with Bonferroni post-doc tests: between groups, F=11.02, $p$<0.0001. |
| Fig S7A | Membrane potential ramps at different time window, during PV+ inactivation. | Same as Figure 6C-D | Including all 23 control recordings (spiking and non-spiking ones); n=11 spiking neurons used in Figure 4C; n=12 non-spiking neurons first used in the figure.<br><br>Including all spiking and non-spiking recordings during PV+ inactivation: 11 out of 14 recordings in Figure 5G-J, which meet the criteria (see methods);<br><br>Including all spiking and non-spiking recordings during SOM+ inactivation: 11 out of 12 recordings in Figure 5G-J, which meet the criteria (see methods). | None | Spearman's correlation:<br>-14~-0s window, r=0.43, $p$=0.0000;<br>-14~-6s window, r=0.41, $p$=0.0001;<br>-6~0s window, r=-0.02, $p$=0.8993. |
| Fig S7B | Membrane potential ramps at different time window, during SOM+ inactivation. | | | | Spearman's correlation:<br>-14~-0s window, r=-0.10, $p$=0.2364;<br>-14~-6s window, r=-0.03, $p$=0.7602;<br>-6~0s window, r=-0.04, $p$=0.7285. |
| Fig S7C | Membrane potential during preparation of movement, during PV+ inactivation. | Superficial recordings during PV+ inactivation across 169~391μm; depth:338±48μm.<br><br>Superficial recordings during SOM+ inactivation across 190~430μm;<br>depth: 359±41μm | Superficial recordings during PV+ inactivation: 7;<br>Control superficial recordings: 12;<br><br>Superficial recordings during SOM+ inactivation: 7. | None | Two-way ANOVA with Bonferroni post-hoc tests; comparision of different groups, F=11.10, $p$<0.0001. |
| Fig S7D | Membrane potential during preparation of movement, during SOM+ inactivation. | | | | two-way ANOVA with Bonferroni post-hoc tests; comparision of different groups, F=0.7720, $p$=0.2454. |

**Supplemental Table 2.**

Synaptic weights in the reduced model of the local PFC circuit.

| ***Presynaptic*** | ***Postsynaptic*** | ***Weight (AU)*** |
|---|---|---|
| External inputs | Principal neuron | 1.0 |
| | PV+ interneuron | 1.0 |
| | SOM+ interneuron | 0.3 |
| Principal neuron | Principal neuron | 1.0 |
| | PV+ interneuron | 0.01 |
| | SOM+ interneuron | 0.01 |
| PV+ interneuron | Principal neuron | –1.0 |
| | PV+ interneuron | –0.25 |
| | SOM+ interneuron | –0.01 |
| SOM+ interneuron | Principal neuron | –0.25 |
| | PV+ interneuron | –1.0 |
| | SOM+ interneuron | 0 |
